# Modulation of the tumour promoting functions of cancer associated fibroblasts by phosphodiesterase type 5 inhibition increases the efficacy of chemotherapy in human preclinical models of esophageal adenocarcinoma

**DOI:** 10.1101/2020.04.21.052647

**Authors:** Annette Hayden, Antigoni Manousopoulou, Andrew Cowie, Robert Walker, Benjamin P. Sharpe, Jack Harrington, Fereshteh Izadi, Ewan Kyle, John H. Saunders, Simon L. Parsons, Alison A. Ritchie, Philip A. Clarke, Pamela Collier, Spiros D. Garbis, Matthew Rose-Zerilli, Anna M. Grabowska, Timothy J. Underwood

## Abstract

**Background and aims:** Esophageal adenocarcinoma (EAC) is chemoresistant in the majority of cases. The tumor-promoting biology of cancer associated fibroblasts (CAF) make them a target for novel therapies. Phosphodiesterase type 5 inhibitors (PDE5i) have been shown to regulate the activated fibroblast phenotype in benign disease. We investigated the potential for CAF modulation in EAC using PDE5i to enhance the efficacy of chemotherapy.

**Methods:** EAC fibroblasts were treated with PDE5i and phenotypic effects examined using immunoblotting, immunohistochemistry, gel contraction, transwell invasion, organotypics, single cell RNAseq and shotgun proteomics. The combination of PDE5i with standard-of-care chemotherapy (Epirubicin, 5-Fluorouracil and Cisplatin) was tested for safety and efficacy in validated near-patient model systems (3D tumor growth assays (3D-TGAs) and patient derived xenograft (PDX) mouse models).

**Results:** PDE5i treatment reduced α–SMA expression in CAFs by 50% (p<0.05), associated with a significant reduction in the ability of CAFs to contract collagen-1 gels and induce cancer cell invasion, (p<0.05). RNAseq and proteomic analysis of CAF and EAC cell lines revealed PDE5i specific regulation of pathways related to fibroblast activation and tumor promotion. 3D-TGA assays confirmed the importance of stromal cells to chemoresistance in EAC, which could be attenuated by PDE5i. Chemotherapy+PDE5i in PDX-bearing mice was safe and significantly reduced PDX tumor volume (p<0.05).

**Conclusion:** PDE5 is a candidate for clinical trials to alter the fibroblast phenotype in esophageal cancer. We demonstrate the specificity of PDE5i for fibroblasts to prevent transdifferentiation and revert the CAF phenotype. Finally, we confirm the efficacy of PDE5i in combination with chemotherapy in close-to-patient *in vitro* and *in vivo* PDX-based model systems.

## Introduction

Esophageal adenocarcinoma (EAC) is usually lethal. Most patients present with late stage disease and for those amenable to potentially curative treatments, 5-year survival is 50% at best. Randomised controlled trials (RCTs) confirm a survival advantage for neoadjuvant chemotherapy +/− radiotherapy, but this benefit is restricted to a minority of patients. For the majority, neoadjuvant treatments are ineffective, significantly morbid and delay definitive surgery^1-3^.

Large-scale genome sequencing studies have detailed the genetic landscape of EAC and identified potential molecular targets. These data reveal a highly complex tumor with driver gene mutations present in non-malignant precursor lesions that never progress to cancer, suggesting that drivers of disease development and progression may lie within the tumor microenvironment^4-7^.

We have previously reported that activated cancer-associated fibroblasts (CAFs) influence outcome in EAC and the biological properties of esophageal CAFs that promote tumor progression^8, 9^. CAFs have also been shown to influence the immune cell infiltrate and response to chemotherapy in a range of tumors^10-12^. In general, the tumor-promoting properties of CAFs have been associated with the alpha-smooth muscle actin (α-SMA)-positive, activated, myofibroblast phenotype observed in cancer, fibrosis and wound healing^13-15^. CAF targeting strategies have mostly focussed on the effectors of CAF tumor promotion including cell signalling and extracellular matrix molecules. We have been working to understand how to target the CAF phenotype itself and whether new or existing drugs can be purposed for this use.

Phosphodiesterase type 5 (PDE5) is part of a complex superfamily of hydrolases ^16^ that control cAMP and cGMP levels by catalysing their breakdown. PDE5 is widely expressed in normal tissue and many human cancers, and its inhibition results in an upregulation of cGMP, which activates several downstream pathways including protein kinase G (PKG) signalling. Downstream substrates of PKG are implicated in a variety of biological processes such as smooth muscle contraction, cell differentiation, proliferation, adhesion and apoptosis^17, 18^. The main function of PDE5 is to control vascular tone by regulating intracellular cGMP and calcium levels (Figure 1).

**Figure 1.**
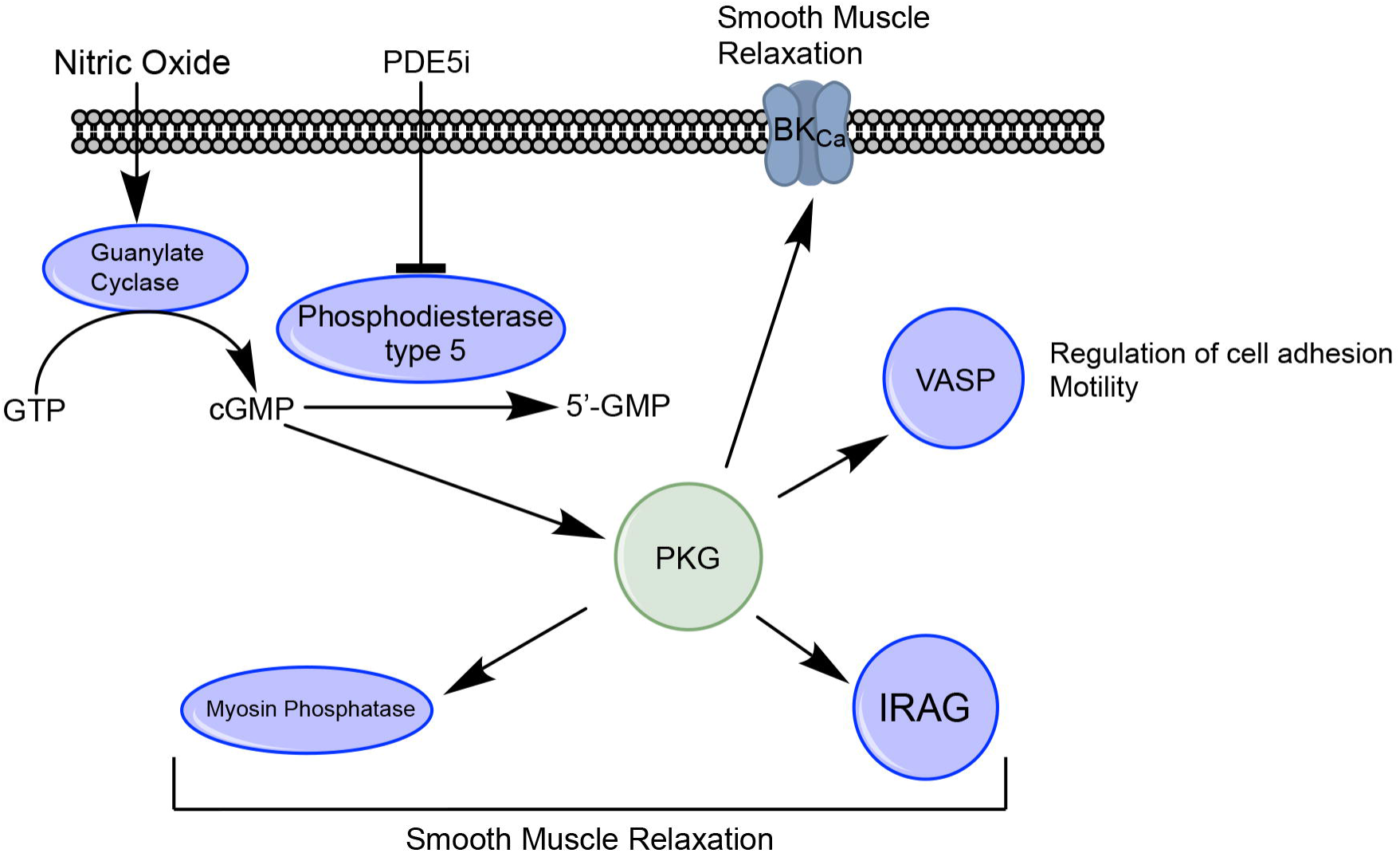
The involvement of PDE5 in the NO/cGMP/PKG pathway. Nitric oxide produced by endothelial cells activates guanylate cyclase in smooth muscle and stimulates cGMP production. cGMP activates downstream signals including protein kinase G family proteins. Targets of PKG in turn stimulate smooth muscle relaxation. VASP is also a major target of PKGs, and regulates cell adhesion and motility.

Phosphodiesterase type 5 inhibitors (PDE5i) were first licensed to treat erectile dysfunction. More recently, high doses have been approved to treat pulmonary arterial hypertension and lower urinary tract symptoms^19-22^. New studies suggest repurposing PDE5i for treating conditions such as cancer or lung disease^18, 23^. PDE5i have been found to attenuate the myofibroblast phenotype of prostatic fibroblasts, suggesting they could target the inflammatory/activated microenvironment observed in many solid tumors^24^.

In EAC, we hypothesized that directly targeting the CAF phenotype with PDE5i would downregulate the tumor-promoting effects of CAFs and improve EAC sensitivity to conventional chemotherapy. This may improve outcomes for patients with EAC, of whom up to 80% do not respond to standard-of-care neoadjuvant treatment^25^. Recent evidence has shown multimodal therapies of this type have acceptable tolerability and therapeutic potential^26-28^.

In this study we characterized PDE5 expression in the human esophagus and described the effect of PDE5is on the tumor-promoting functions of esophageal CAFs in 2D and 3D models *in vitro*. We documented changes in CAF protein expression in response to PDE5i using shotgun proteomics and applied single cell RNA sequencing to demonstrate a phenotypic change in CAFs driven by co-culture with cancer cells and inhibited by PDE5i treatment. Finally, we moved to a validated near-patient EAC model system to assess tolerability and efficacy of PDE5i in combination with standard-of-care chemotherapy.

## Materials and Methods

### Tissue and cell collection, maintenance and RNAi

Tissue was collected and stored with ethical agreement and informed consent (REC: 09/H0504/66 and 18/NE/0234) at University Hospital Southampton, and fibroblasts were extracted from normal esophagus and esophageal adenocarcinoma and sub-cultured as previously described^29^. siRNA-mediated silencing of PDE5 was performed using INTERFERin transfection reagent and two commercially available siRNA sequences (detailed in supplementary methods). Endoscopic tumour biopsies (REC: 10/H0401/80) and fresh surgical specimens (REC: 08/H0403/37) were collected with informed consent from patients at Nottingham University Hospitals NHS Trust in 2014-15, and used in accordance with National Research Ethics Service approval to generate oesophageal cancer cell-lines (detailed in supplementary methods) as previously described^30^.

### Immunohistochemistry

Optimization and staining of cohort on full face sections using polyclonal rabbit anti-PDE5 (ab224232, abcam) was performed on a Dako link automated staining machine according to the manufacturer’s instructions.

### Western blotting

Primary normal esophageal fibroblasts were treated under different conditions: ± TGF-β1, ± vardenafil (V-902, Sigma), and TGF-β1 + vardenafil; cancer associated fibroblasts were treated with vardenafil, or vehicle. Cells were lysed and clarified after 72 hours. For full details of antibodies used, experimental conditions and western blotting see supplementary methods.

### Immunofluorescence

Immunofluorescence analyses were carried out as previously described^31^. Primary antibodies: rabbit polyclonal anti-PDE5 (ab224232, abcam) and mouse monoclonal anti-α-SMA (M085129-2, Dako), Secondary antibodies: Alexa Fluor 568 donkey anti mouse IgG and Alexa Fluor 488 goat anti rabbit IgG (Molecular Probes). Cell nuclei were counterstained with 1 μg/ml DAPI (4’,6’-diamidino-2-phenylindole).

### Gel contraction

Gel contraction assays were conducted as previously described^15^. Briefly, 0.5×10^6^ NOFs or CAFs (± TGF-β1, +/− 72 hours vardenafil treatment) were seeded in collagen-1 gels. Gels were photographed and weighed after 24 hours.

### Transwell invasion

Transwell invasion assays were conducted as previously described^32^, using conditioned medium from NOFs or CAFs as the chemoattractant. NOFs and CAFs were cultured in the presence of TGF-β1 or TGF-β1 + vardenafil for 72 hours before washing out and collecting conditioned medium after 24 hours. FLO-1 cells were plated in the top chamber in identical numbers. All experiments were repeated 3 times with 4 replicates per experiment. Normalization was performed to the mean value of replicate 1 for all experimental conditions and data expressed as % invasion compared to the vehicle control.

### Organotypic cultures

Organotypic cultures were carried out as previously described^29, 32^. Briefly, CAFs were pre-treated with 50μM vardenafil or vehicle control for 72 hours before plating in 1:1 collagen:matrigel gels at a density of 5×10^5^ cells per gel, and the next day 5×10^5^ FLO-1 cells were cultured on top of the gels. Organotypic cultures were raised onto steel grids with overlaid nylon membrane supports and cultured at the air-liquid interface overnight before treatment. Organotypic cultures were incubated for a further ten days with added Vardenafil or vehicle control before processing. Invasion was quantified and compared as previously described^33^.

### Proteomics

#### Quantitative proteomics analysis

Cells were treated with vardenafil, vehicle or PDE5 siRNA for 72 hours before trypsinisation and clarification. Cells were lysed and protein extracts were quantified. One hundred ug of protein per sample was reduced, alkylated and enzymatically digested using trypsin. Peptides were labelled using the isobaric tag for relative and absolute quantitation (iTRAQ) 8-plex reagents and analysed using two-dimensional liquid chromatography and tandem mass spectrometry as reported previously^34, 35^. For full details of downstream processing see supplementary methods.

### Droplet barcoded single cell RNA sequencing

CAFs were grown in isolation or in co-culture MFD-1 cell line^36^. Cells were treated with vardenafil for 72 hours before analysis. This model was used to look at the transcriptional regulation of both cancer cells and CAFs in the presence of PDE5 inhibition.

Cultured cells were trypsinized and resuspended in a cell suspension buffer and single cell RNA seq libraries created using DropSeq and then sequenced using Illumina NextSeq500. Single cell RNA-seq data was processed in R version 3.5.1 (Feather Spray) to cluster cells and perform differential gene expression analysis. For full details of downstream processing see supplementary methods.

### 3D-tumor growth assay

The individual patient’s epithelial cells were established, characterised, and then co-cultured in the 3D-TGA +/-human mesenchymal stem cells (hMSC)^30^. Following exposure to drug combinations at human tissue-relevant concentrations, the cell viability was assessed using the alamarBlue® assay (ThermoFisher Scientific, Loughborough, UK). Viability curves were generated and IC_50_ values calculated using GraphPad Prism 5 software (San Diego, CA, USA). These were compared with the mean peak serum concentration achieved in patients for each chemotherapeutic agent to evaluate chemotherapeutic response as previously reported^30^. For full details see supplementary methods.

### Patient-derived xenograft models

Patient-derived xenografts were developed using EAC biopsy specimens directly implanted into immuno-deficient mice as previously described^37^. Due to loss of the human stromal compartment in such models, human mesenchymal stem cells (hMSCs) were co-implanted with the xenograft and supplemented before treatment began. Mice were treated with vehicle, Epirubicin, Cisplatin and Capecitabine (ECX) or Cisplatin and Capecitabine (CX) alone or in combination with vardenafil or tadalafil. For full details see supplementary methods.

### Statistical analysis

Statistical analysis was performed with SPSS® version 19 (SPSS, Chicago, United States). Patient survival was plotted using the Kaplan-Meier method and analyzed using the Log-rank test, with patients censored at last follow-up. Multivariate Cox logistic regression was used to assess the relationship between α-SMA and pathological outcomes after resection. Kruskal-Wallis, Mann Whitney *U* and T-tests were used to compare groups, as appropriate. *P* < 0.05 was considered statistically significant (P < 0.05*, P <0.01**).

## Results

### Characterisation of PDE5 inhibitors in esophageal cancer

To assess the suitability of PDE5 as a novel target in EAC we determined the expression of PDE5 in esophageal cancer cells and CAFs in primary resected tumor tissue and in extracted primary cells. PDE5 was highly and ubiquitously expressed in esophageal cancer tissue with low expression in normal tissue (Figure 2A). In matched normal esophageal fibroblasts (NOFs) and CAFs, 2 commonly used EAC cell lines and a primary epithelial esophageal cancer cell line extracted in our laboratory (MFD-1)^36^, variable PDE5 expression was observed, determined by cell sub-type. There was little or no PDE5 protein expression in the cancer cell lines with highest expression in CAFs (Figure 2B). Although a heterogeneous population, in general, CAFs are associated with a contractile and secretory phenotype characterized by increased expression of alpha-smooth muscle actin (αSMA). This activated, myofibroblastic state in cancer is believed to be driven by cancer cell signaling and can be recapitulated by treating normal fibroblasts with TGF-β1 *in vitro*. We repeated our previous experiments to confirm that NOFs treated with TGF-β1 significantly induced αSMA expression^8^, but when co-treated with vardenafil (a specific PDE5i) the increase in αSMA expression was abolished (Figure 2C). Having established the potential of PDE5i to prevent NOF trans-differentiation *in vitro* we explored the possibility that PDE5i could suppress αSMA expression in CAFs. After 72 hours of culture with vardenafil CAFs reduced αSMA expression by over 50% (p<0.01; Figure 2D). The on-target effects of PDE5i were confirmed by observing appropriate decreases in PDE5 and αSMA protein expression in response to PDE5 siRNA (Supplementary Figure S1). Importantly for potential *in vivo* applications, we found that daily dosing of PDE5i produced significant αSMA downregulation compared to a single application of PDE5i-containing medium 72 hours before analysis (Supplementary Figure S2A). After withdrawal of PDE5i, αSMA expression returned to pre-treatment levels within 72 hours (Supplementary Figure S2B).

**Figure 2.**
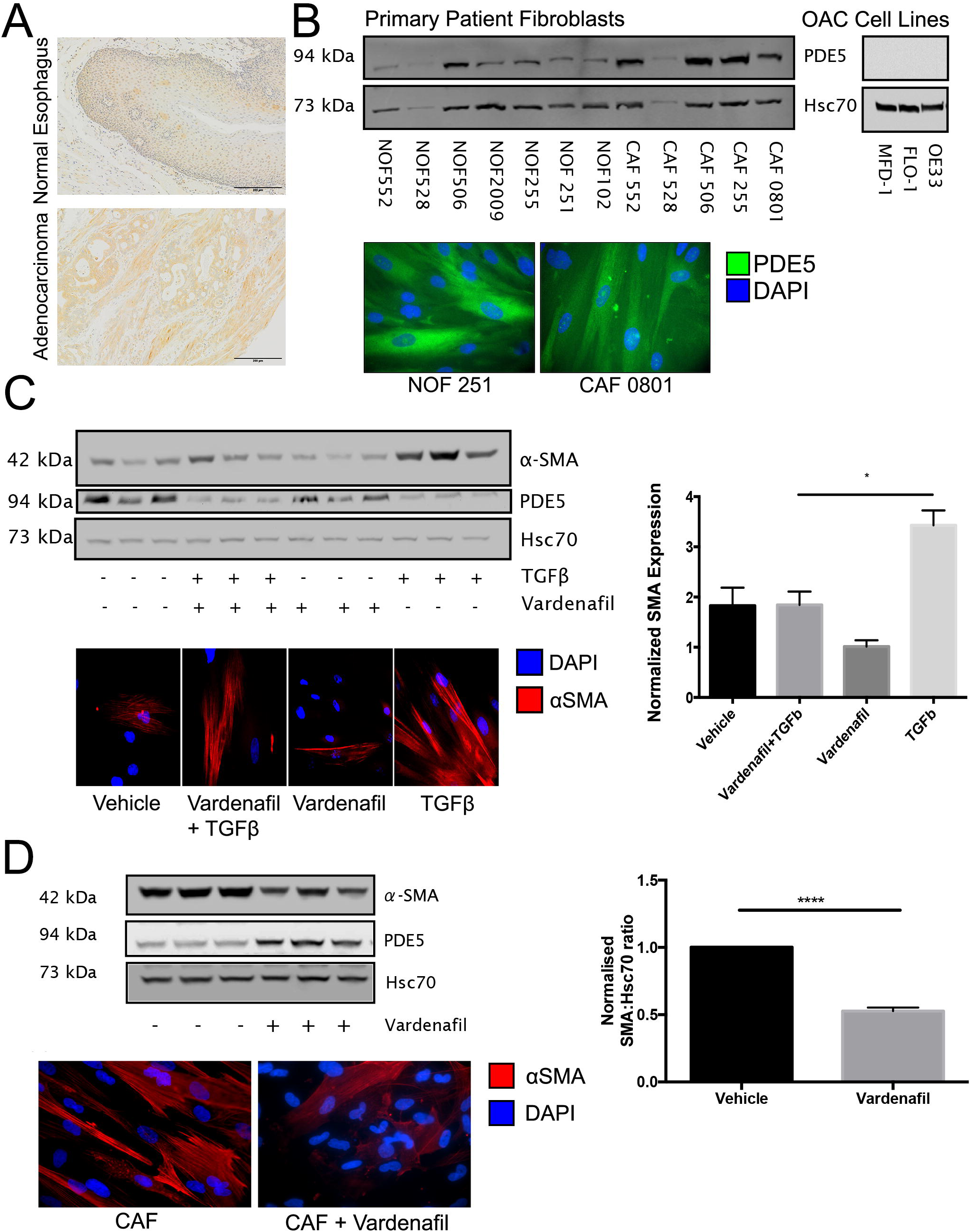
**A**. PDE5 expression analysed by IHC of normal esophagus and esophageal adenocarcinoma. PDE5 expression is highest in but not confined to the stromal tissue. **B**. Western blot and ICC for PDE5 expression in three cancer cell lines and five matched normal (NOF)/cancer (CAF) primary esophageal fibroblasts. **C**. NOFs treated with TGF-β1, +/-50μM vardenafil for 72 hours. TGF-β1 treated NOFs express higher α-SMA, vardenafil pre-treatment abrogated the TGF-β1 induced expression of α-SMA. **D**. CAFs treated with 50μM vardenafil for 72 hours reduced α-SMA expression by 50% by western blot and ICC.

### PDE5 inhibition reduces fibroblast contraction and esophageal cancer cell invasion

The expression of αSMA is characteristic of the myofibroblast phenotype but does not necessarily indicate functional capacity. To test the hypothesis that PDE5i treatment of myofibroblasts could interfere with known tumor-promoting functions we performed a series of *in vitro* experiments to assess extracellular matrix contraction and the promotion of cancer cell invasion. NOFs were embedded in collagen-1 gels after being treated with TGF-β1 alone or TGF-β1 + vardenafil. Fibroblasts that were treated with TGF-β1 significantly increased both αSMA expression and collagen-1 gel contraction, but after pretreatment with vardenafil the induction of αSMA and gel contraction was substantially and consistently reduced (Figure 3A). CAFs express high levels of αSMA and induce collagen-1 gel contraction. Treatment with vardenafil significantly reduced αSMA expression and collagen-1 gel contraction in CAFs (Figure 3B). Next, we assessed the ability of conditioned medium taken from fibroblast cultures to promote cancer cell invasion in transwell invasion assays under a variety of conditions. The conditioned medium from TGF-β1-treated NOFs promoted 5 times more invasion of esophageal adenocarcinoma cells than conditioned medium from vehicle-treated NOFs, whereas when vardenafil was added to TGF-β1 treatment of NOFs the resultant conditioned medium did not promote invasion (Figure 3C). Similarly, vardenafil-treated CAF-conditioned medium induced significantly less cancer cell invasion than vehicle-treated CAF-conditioned medium (Figure 3D). Similar observations were made using PDE5 siRNA (Supplementary Figure S1). This finding was reproduced in the more physiologically relevant organotypic co-culture model where we observed that vardenafil-treated CAFs had lost their ability to promote cancer cell invasion, compared to vehicle treated CAF (Figure 3E). These findings suggested that, *in vitro*, PDE5i treatment was able to suppress both the transdifferentiation of NOFs and the tumor-promoting characteristics of CAF that we have previously observed^8^.

**Figure 3.**
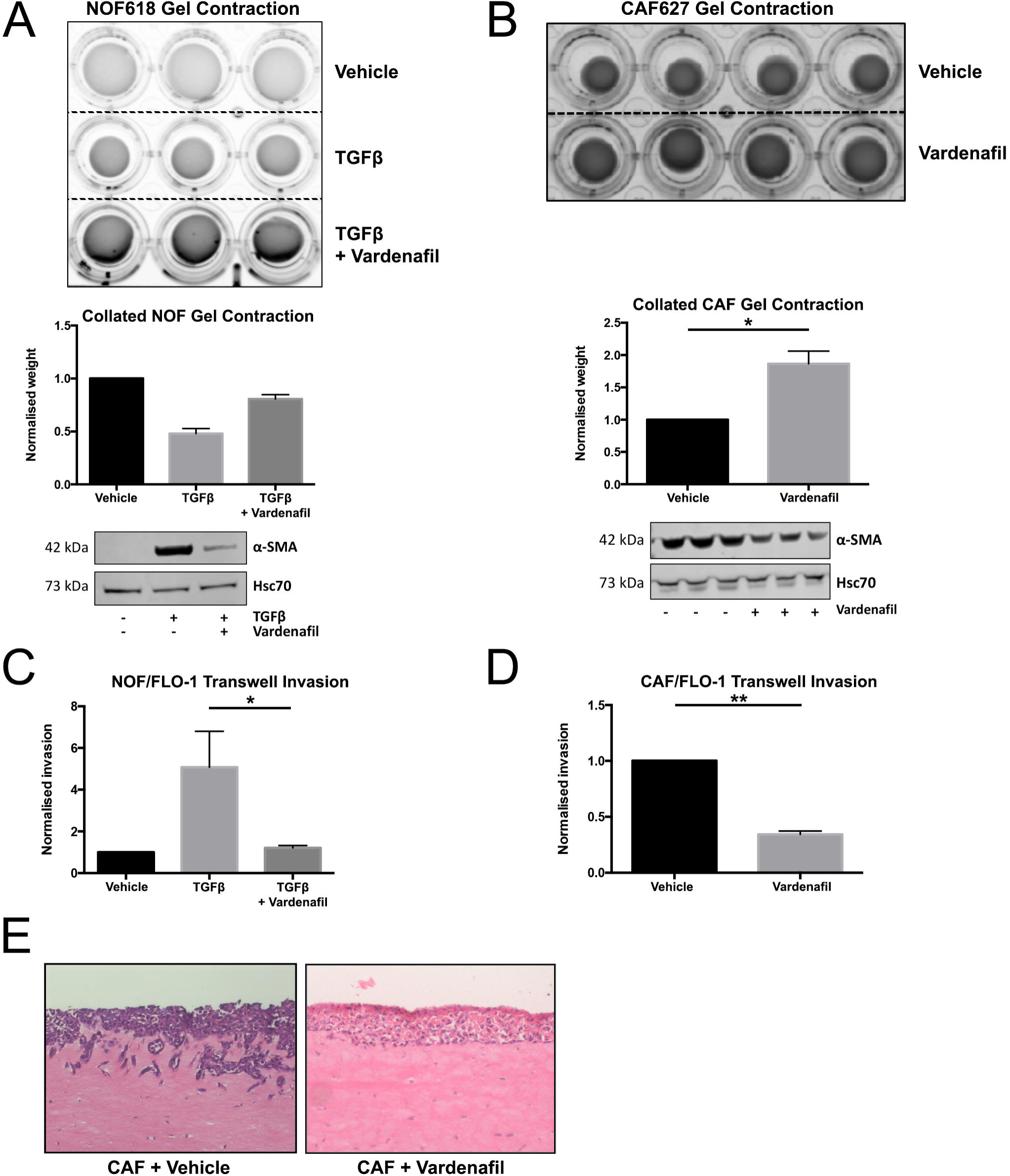
**A.** Fibroblast contraction analysed by normal esophageal fibroblasts (NOFs) treated with TGF-β1 ± 50μM vardenafil embedded in collagen-1 gel. TGF-β1 treatment induced αSMA expression and gel contraction in normal fibroblasts, co-treatment with vardenafil inhibited the upregulation of αSMA and reduced gel contraction by 50%. **B**. Cancer associated fibroblast (CAF) contraction was analysed by collagen-1 gel contraction, ± 50μM vardenafil. αSMA expression and collagen-1 gel contraction is 2-fold greater in untreated CAFs compared to vardenafil treated CAFs. **C,D** Cancer cell invasion was analysed by transwell assays using conditioned medium from fibroblasts as the chemoattractant. **C**. Normal fibroblasts ± TGF-β1 ± vardenafil, TGF-β1 treated NOFs promoted invasion of FLO-1 cells with a 5-fold induction. This induction was abrogated by vardenafil treatment. **D**. CAFs treated with vardenafil reduced invasion of FLO-1 cells by 60%. **E**. Organotypic co-culture of FLO-1 cells and CAFs also showed inhibition of invasion when treated with vardenafil.

### Proteomic analysis of fibroblasts treated with vardenafil or PDE5 siRNA identifies modulation of major pathways associated with the cancer promotion

In keeping with previous reports in benign disease^24^ we established the ability of PDE5i to ameliorate some of the tumor-promoting functions of TGF-β1-driven, activated esophageal fibroblasts *in vitro*. To explore the cellular events responsible for these effects we took a whole-proteome based approach. We have previously demonstrated the benefits of this approach to identify pathways and participating proteins that may provide novel insight into the tumor-promoting properties of CAFs^34^.

Proteomic analysis was carried out on a NOF/CAF patient-matched pair. The CAFs were treated with vehicle (negative control), vardenafil (PDE5i), PDE5 siRNA (positive control) and negative control siRNA and total protein expression assessed by quantitative proteomic profiling. To examine the effects of vardenafil or PDE5i siRNA treatment on the global proteomic profile of CAFs, we considered the following log2ratios: PDE5i vs. CAF vehicle, PDE5 siRNA vs. siRNA negative control and CAF vehicle vs. NOF. In total, 8118 proteins were quantified across all analysed samples (peptide level FDR < 0.05) **(Supplementary Table x1)**. Principal component analysis of all quantified proteins showed that vardenafil treated CAFs clustered together with PDE5 siRNA treated CAF compared to vehicle treated CAF **(Figure 4A)**. Since their global proteomic profile was similar, we considered PDE5i vs. CAF vehicle and PDE5 siRNA vs. siRNA negative control as one group and performed a one-sample T-Test to identify differentially expressed proteins (DEPs) following treatment with vardenafil or PDE5 siRNA. In total, 812 proteins were found to be up-regulated and 725 down-regulated in CAFs treated with vardenafil or PDE5 siRNA compared to their respective controls (**Supplementary Table x2)**. In order to identify which of these proteins reflect the amelioration of the CAF phenotype following vardenafil or PDE5 siRNA treatment, we compared the DEPs in CAFs treated with vardenafil or PDE5 siRNA to our previously published dataset of DEPs in CAFs vs. NOFs^34^. Using this approach, we identified 83 proteins that were down-regulated in CAFs vs. NOFs but became up-regulated in CAFs following treatment with vardenafil or PDE5 siRNA **(Figure 4B)**. Conversely, we identified 88 proteins that are up-regulated in CAFs vs. NOFs but became down-regulated in CAFs following treatment with vardenafil or PDE5 siRNA **(Figure 4B) (Supplementary Table x3**). We then performed gene ontology analysis for these 171 proteins that reversed their trend of modulation following treatment with vardenafil or PDE5 siRNA compared to CAFs. ECM organization (p = 0.01), ECM disassembly (p = 0.0008), sequestering of TGFβ in ECM (p = 0.0003), regulation of extracellular exosome assembly (p = 0.0005), cell-cell adhesion (p = 0.01), cell migration (p = 0.02), DNA damage response (p = 0.003), regulation of apoptosis (p < 0.0001), programmed cell death (p = 0.002), angiogenesis (p = 0.02), response to hypoxia (p = 0.02) and insulin receptor signaling pathway (p = 0.006) were significantly over-represented GO terms **(Figure 4C)**. These are all major pathways associated with the cancer-promoting properties of fibroblasts and identified in our previous studies of EAC fibroblasts^34^. To provide additional granularity, proteins exhibiting the most significant changes in expression in response to PDE5i treatment have been represented on a heatmap with the corresponding gene ontology term highlighted **(Figure 4D)**.

**Figure 4.**
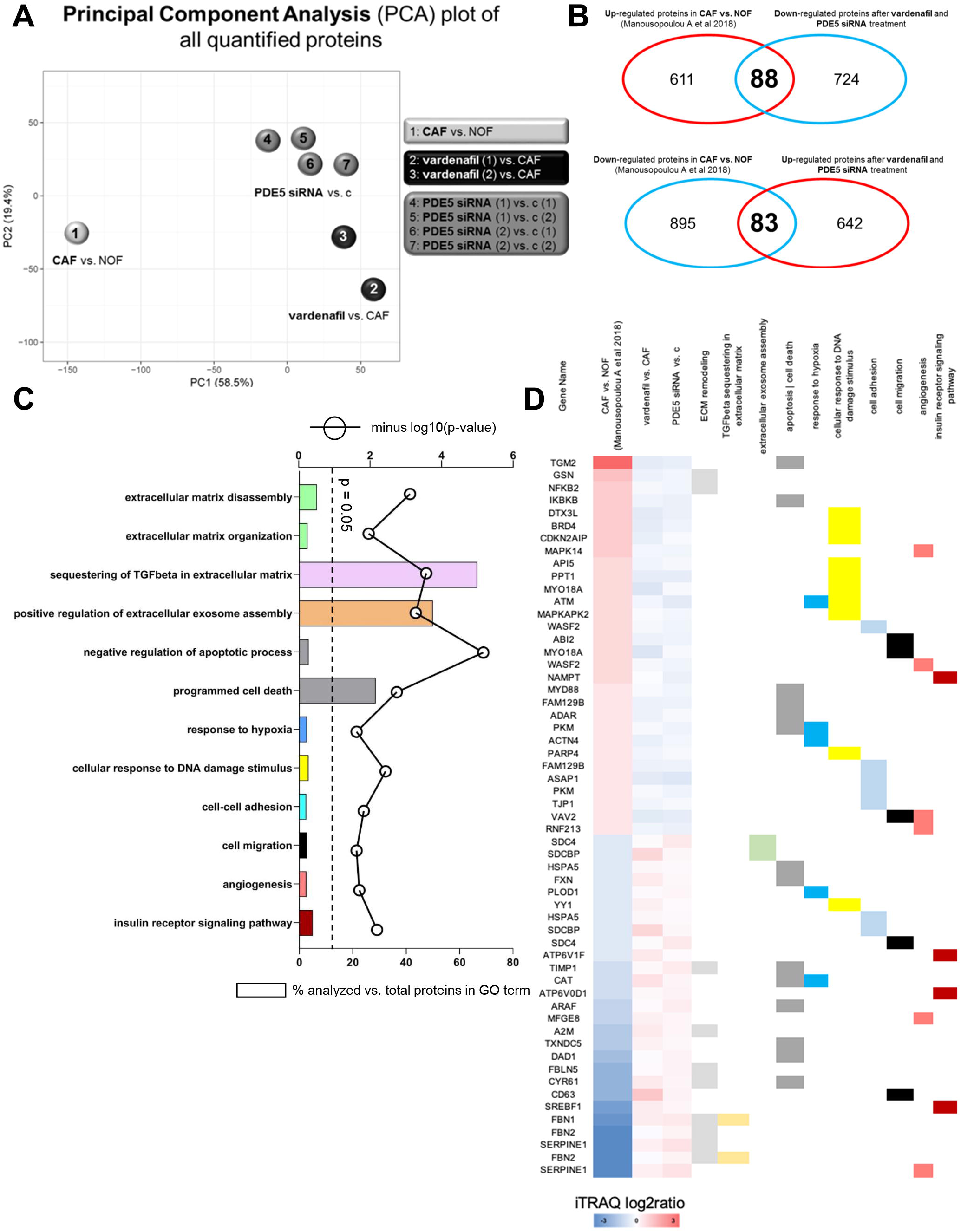
Proteomics: **A**. Principal component analysis of all quantified proteins showed that vardenafil treated CAFs clustered together with PDE5 siRNA treated CAF compared to vehicle treated CAF. **B**. Comparison of DEPs in CAFs treated with vardenafil or PDE5 siRNA with a previously published dataset of DEPs in CAFs vs. NOFs^34^, identified 83 proteins down-regulated in CAFs vs. NOFs but became up-regulated in CAFs following treatment with vardenafil or PDE5 siRNA. Conversely, 88 proteins were up-regulated in CAFs vs. NOFs but became down-regulated in CAFs following treatment with vardenafil or PDE5 siRNA. **C**. Gene ontology analysis using DAVID of the 171 proteins that reverse their trend of modulation following treatment with vardenafil or PDE5 siRNA compared to CAFs. **D**. Proteins mapping to the respective GO terms.

In summary, these findings suggest that vardenafil is specific for PDE5 inhibition in *ex-vivo* esophageal CAFs and leads to down-regulation of established cancer-promoting CAF pathways.

### Single cell RNA sequencing reveals suppression of activated CAF phenotypes in MFD-1/CAF co-cultures treated with a PDE5 inhibitor

To this point, experiments had focused on understanding the specificity of PDE5i in prevention of fibroblast trans-differentiation and the functional/phenotypic effects of PDE5i on CAFs. To be useful as a potential CAF-targeting treatment in cancer, these effects would need to be retained in the presence of cancer cells and be able to overcome any cancer cell-derived CAF-promoting signaling. To explore this, we took a single cell whole-transcriptomic approach using droplet-based microfluidics and single cell RNA sequencing (DropSeq) to analyse the gene expression of individual CAFs and esophageal cancer cells in direct co-culture^38^. CAFs were grown in isolation or in co-culture with esophageal cancer cell line, MFD-1 ^36^. Cells were treated with vardenafil for 72 hours before analysis. This model was used to look at the transcriptional regulation of both cancer cells and CAFs in the presence of PDE5 inhibition.

Unsupervised clustering produced 2 broad clusters of cells, identified as either MFD1 (cancer cells) or CAFs, as defined by their transcriptomic profiles **(Figure 5A)**. Within these clusters a further 8 sub-clusters were identified **(Figure 5B)**. In general, the cells clustered based on their cell type and within those clusters on their culture conditions.

**Figure 5.**
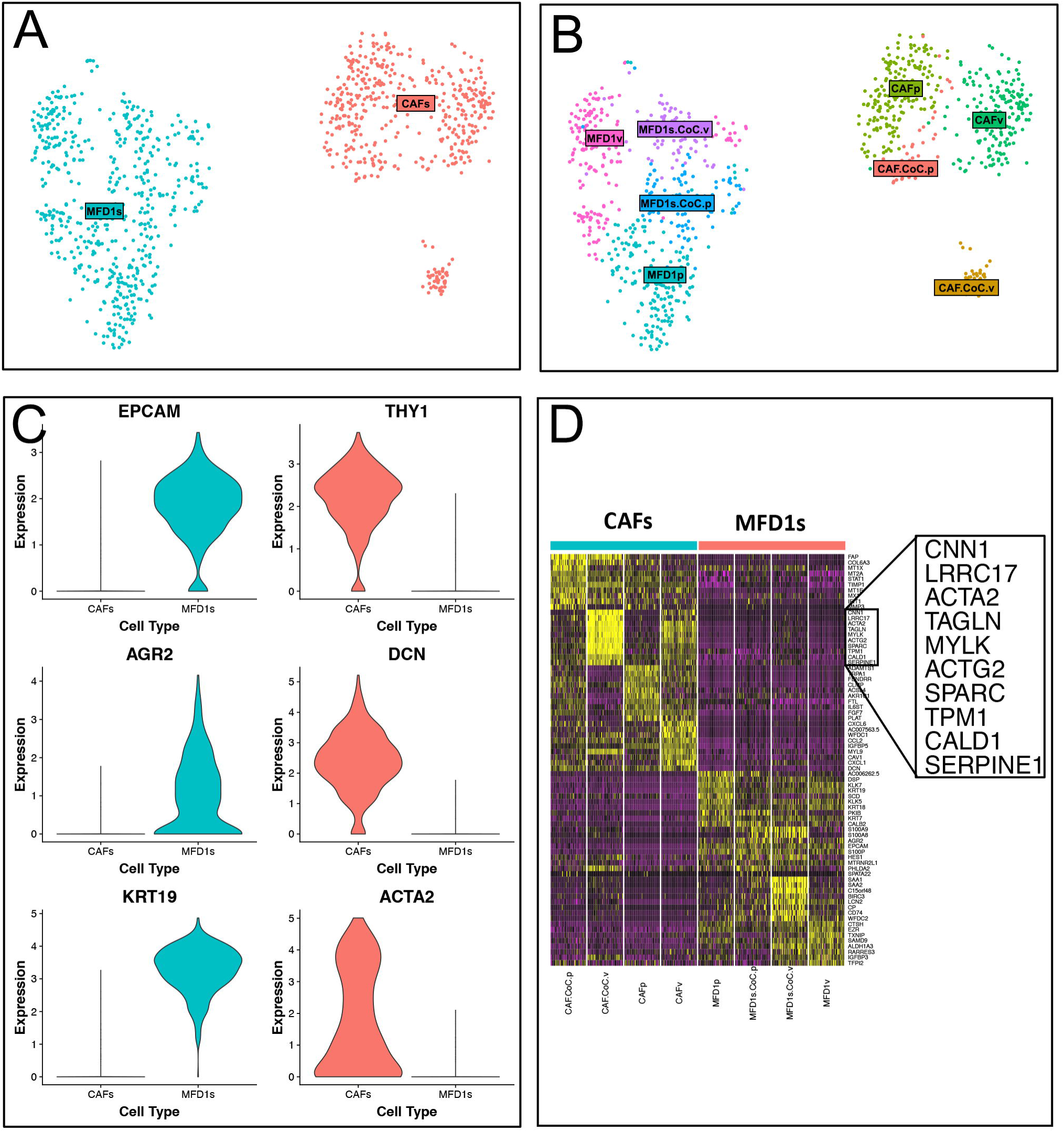
Droplet barcoded single cell RNA sequencing. **A**. Cluster analysis of MFD-1 and primary CAFs based on differentially expressed genes from single cell RNA sequencing analysis. Firstly cells were clustered as either cancer cells (MFD-1) (blue) or fibroblasts (red). **B**. Further cluster analysis showed CAFs treated with PDE5 inhibitor vardenafil (CAFp) clustered separately from vehicle-treated CAFs (CAFv). PDE5i-treated co-cultured CAFs clustered with all other CAF cultures, whereas vehicle-treated CAF co-cultured with MFD-1 show a phenotypic shift and clustered separately. **C**. Violin plots showing typical gene expression profiles for fibroblasts (THY1, DCN, ACTA2) or epithelial cells (KRT8, KRT18, EPCAM) show good separation between the clusters in A. **D**. Heatmap shows the top 5 differentially expressed genes between treatment groups. CAFs and MFD-1 cells have the most distinct gene expression profiles and although PDE5i treatment and co-culture both affect gene expression of both cell types, CAFs were most affected under both conditions.

In order to characterise the individual sub-clusters of cell phenotypes, differential gene expression analysis was performed using Seurat’s FindAllMarkers function with a log fold change cut-off of 1 and otherwise default settings. Canonical marker genes differentially expressed between populations of CAFs (Thy-1/CD90, Decorin (DCN), Smooth Muscle Actin (ACTA2)) and MFD1s (Epithelial cell adhesion molecule (EpCam), Anterior gradient protein 2 homolog (AGR-2), Keratin-19 (KRT19)) are shown in **Figure 5C**.

The most striking finding from this experiment was the differential expression pattern of CAFs grown in different culture conditions. All CAFs from monoculture were similar in their transcriptomic profile to each other (Figure 5B, CAFp & CAFv). Importantly, the PDE5i-treated CAFs from monoculture (CAFp) and co-culture (CAF.CoC.p) clustered together whereas the CAFs co-cultured with MFD1 in the absence of PDE5i formed a distinct cluster (**Figure 5B**, CAF.CoC.v). This cluster was markedly different to those from the monoculture with vehicle treatment and all other CAFs indicating a phenotypic change in CAFs driven by co-culture with cancer cells and inhibited by PDE5i treatment.

The gene expression that defined the transcriptome of CAFs in co-culture is highlighted in Figure 5D. All are associated with the activated/myofibroblast phenotype of CAFs. This data demonstrates that CAFs adopt a myofibroblastic phenotype in co-culture with cancer cells *in vitro* and that this process can be inhibited by treatment with PDE5i, despite the presence of cancer cells, resulting in downregulation of myofibroblast genes such as ACTA2 (αSMA), myosin light chain kappa (MYLK), osteonectin (SPARC) and transgelin (TAGLN)**(Figure 5D)**.

### Three-dimensional co-culture models of close-to-patient cancer cells and human mesenchymal stem cells reveals that PDE5 inhibition increases the efficacy of chemotherapy in EAC

The standard-of-care for locally advanced esophageal adenocarcinoma treatment includes neoadjuvant chemotherapy before resection. Cancer is confirmed by pre-treatment biopsy and this tissue can be used to grow the patient’s cancer epithelial cells *ex vivo*^30^. We have used cancer cells harvested at this time point, as well as matched cells established from post-neoadjuvant chemotherapy surgical specimens, to investigate the potential for PDE5i treatment to enhance the efficacy of standard-of-care chemotherapy (epirubicin, cisplatin and 5-FU; ECF). *Ex vivo* cancer cells (confirmed by FACS) were grown with human mesenchymal stem cells (hMSC) in a 3D matrix to form a 3D-Tumor Growth Assay (3D-TGA) as previously described^30^ **(Figure 6A,B)**. We have previously shown the importance of the stromal compartment in 3D-TGA as the *ex vivo* drug response in the presence, but not absence, of mesenchymal cells accurately reflected clinical chemo-sensitivity, as measured by tumor regression grade (TRG)^30^.

**Figure 6.**
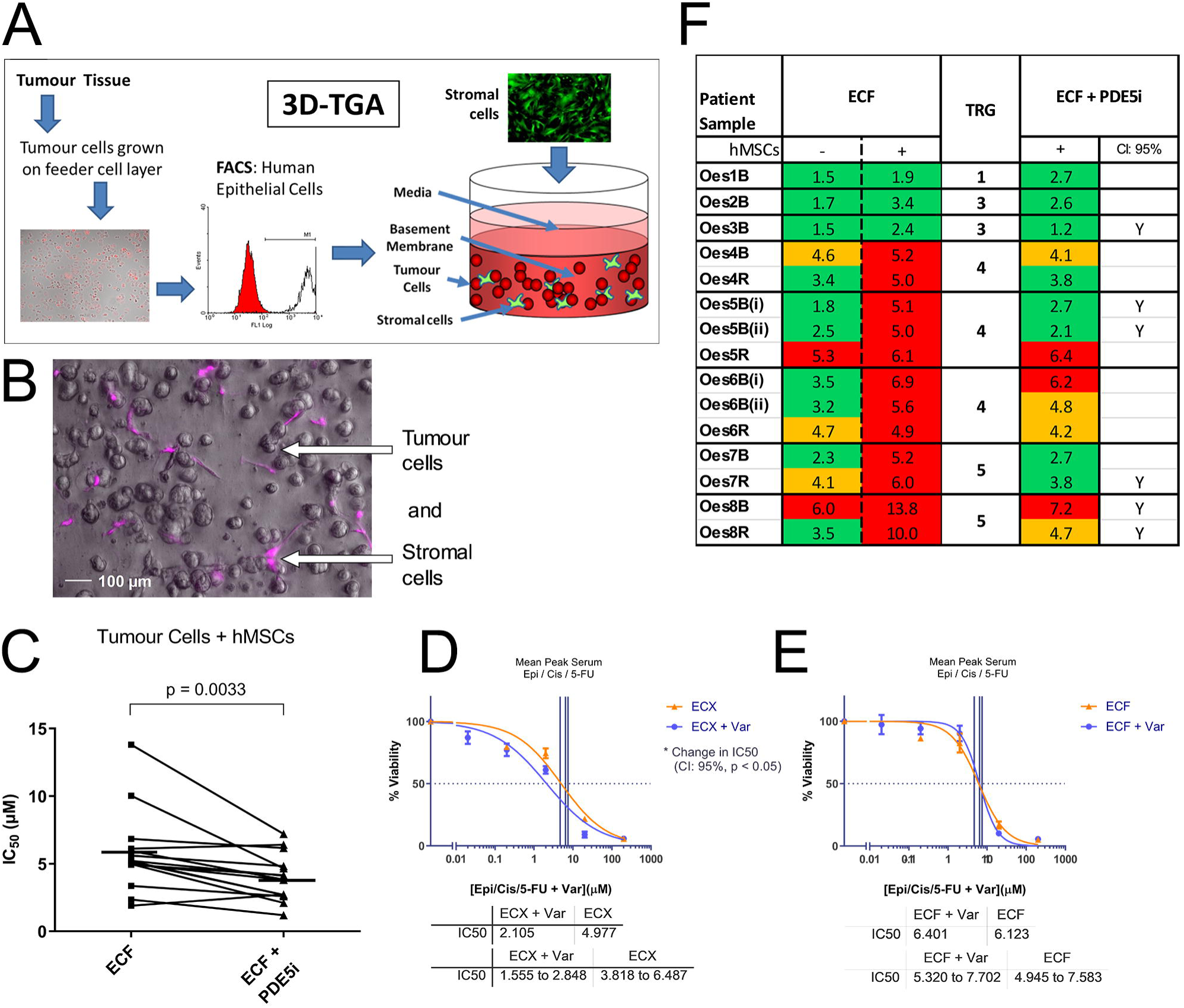
Sensitivity of close-to-patient cells was determined in 3D-TGA, with and without mesenchymal cell co-culture, after 4 day exposure to ECF and Vardenafil (PDE5i) drug combinations. **A**. 3D-TGA workflow. **B**. Image of typical 3D-TGA co-cultured clusters of epithelial cells and mCherry labelled hMSC cells captured using condensed multilayer focussed fluorescent microscopy. **C**. Viability curves were generated and IC_50_ values determined. Overall sensitivity of all the EAC patient samples co-cultured with hMSCs was determined for assays with and without the addition of PDE5i to ECF chemotherapy. Horizontal lines represent mean IC_50_s. Representative patient responses to PDE5i are shown: following the addition of PDE5i to the ECF chemotherapy there is a significant change in chemosensitivity for patient sample Oes5B **D**. but not for Oes5R **E**. Statistical significance is determined by the arithmetic variance between the calculated IC_50_s and their distinct 95% confidence intervals which have no overlap. **F**. IC_50_s obtained for a cohort of EAC patients’ ECF-treated 3D-TGAs, with (+) and without (-) hMSC support and the addition of PDE5i. The patient cancer cell clusters were classified as sensitive (green), borderline (orange), or resistant (red) by comparison of IC_50_ values to the mean peak serum concentrations achieved in patients at the doses used in UK clinical practice. This is marked ‘Y’ where the IC_50_ drop is significant (CI >95%, p <0.005). Tumour regression score (TRG) denotes the chemotherapy response of the patient’s tumour clinically (TRG1-3: sensitive; 4-5: non-responsive).

**Figure 7.**
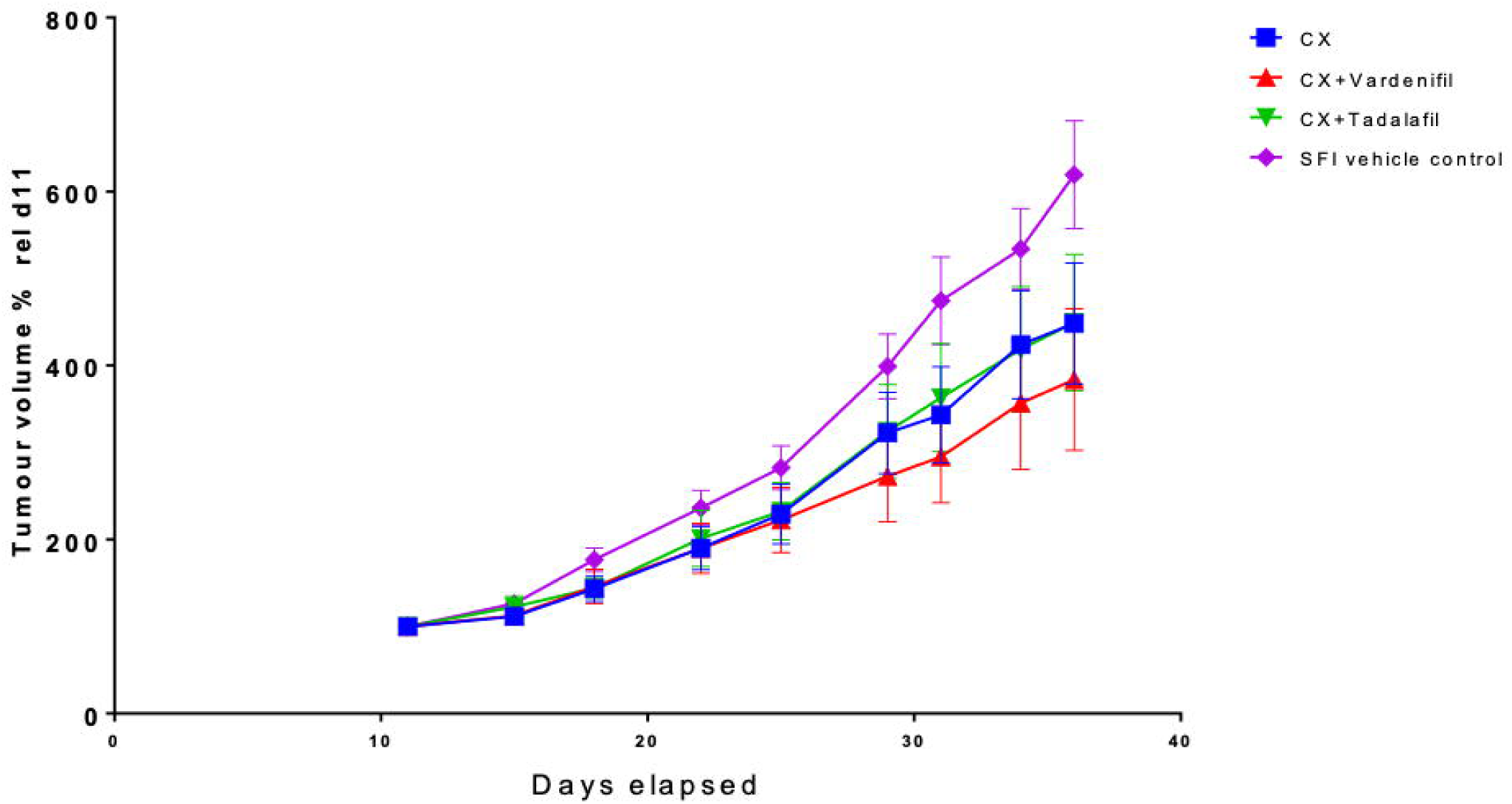
Efficacy in an oesophageal PDX model. An efficacy study was carried out in mice bearing oesophageal PDXs supplemented with human MSCs. Mice were treated with CX, CX+Vard, CX+Tad or vehicle alone. Tumour volume was expressed as a percentage of the last tumour read before commencement of dosing for individual mice and shown as mean +/-standard error of each group.

Using the 3D-TGA we tested the effect of PDE5i on tumor cell chemo-sensitivity in combination with ECF chemotherapy using 15 samples from 8 different patients. We hypothesised that the addition of the PDE5i may affect the interaction of the stromal cells with the epithelial cancer cells within the 3D-TGA, and may reduce the patient-specific ECF chemo-resistance demonstrated by these *ex vivo* 3D co-culture models. When the 3D-TGA models of EAC were exposed to PDE5i alone, there was no chemo-toxic effect seen in any of the 15 models, with or without the hMSC stromal support. PDE5 treatment did not have toxic effects on hMSCs when grown alone in the 3D-TGA **(Supplementary material 3, Figure 1)**.

Overall, the addition of PDE5i to ECF chemotherapy resulted in a significant decrease in IC_50_ chemo-resistance (p=0.0033) (Figure 6C). This effect was not seen without hMSC stromal support **(Supplementary material 3, Figure 2)**, indicating that PDE5i-enhanced chemo-sensitivity is a stromal-mediated process. The individual patient stromal-mediated effect of PDE5i on ECF drug resistance was demonstrated in detail **(Supplementary material 3, Figure 3, 4)**.

A PDE5i-mediated reduction in chemo-resistance was found to be patient-specific **(Figure 6D,E)** with the size of the PDE5i chemo-sensitising effect varying between patients; there was a trend to reduction in IC_50_ seen in 12 of the 15 samples, and a statistically significant (CI >95%, p < 0.05) reduction in the IC_50_ chemo-resistance in 6 of these samples **(Figure 6F)**.

Overall, four of the eight patients studied had a PDE5i mediated significant reduction in ECF chemo-resistance in at least one tumour sample. Five of the eight patients had chemo-resistant tumours, both *ex vivo* in the 3D-TGA and clinically (TRG 4 – 5). With addition of PDE5i to the ECF chemotherapy regimen three of these five patients had a significant reduction in the *ex vivo* IC_50_ to a magnitude similar or less than the mean peak serum concentration used in UK clinical practice, and thus suggesting that 60% of these ECF chemo-resistant tumours could become chemo-sensitive clinically with the adjunctive administration of PDE5i **(Figure 6F)**.

### PDE5i is safe and effective in combination with standard of care chemotherapy in esophageal patient-derived xenograft (PDX) bearing mice

Before considering a human trial of PDE5i in EAC we performed a dose-escalation study to assess potential serious toxicity of combining chemotherapy with PDE5i, and an efficacy study in a PDX mouse model supplemented with human stromal support. The PDX was developed with esophageal cancer tissue taken from patient tumour sample Oes7R. This was chosen as it was resistant to chemotherapy clinically and in the 3D-TGA, but was responsive to adjunctive PDE5i. Since human stroma is lost in PDX models over time, hMSCs were incorporated at passage to maintain a human stroma. A dose escalation study was initially performed with Epirubicin, Cisplatin and Capecitabine (the oral equivalent of 5-FU) (ECX) in three non-tumor bearing mice. This revealed peak tolerable doses of ECX that were 50% of the human equivalent doses (Supplementary Figure S4). Next, dose escalation studies for PDE5i were carried out in four groups of three PDX-bearing mice (no treatment, ECX alone, ECX+PDE5i (vardenafil), ECX+ PDE5i (tadalafil)). PDE5i dose increases were carried out in three phases in combination with a static dose of ECX (as previously determined). No adverse side effects (e.g. weight loss or lowered threshold of ECX tolerability) were reported for maximum doses of either vardenafil or tadalafil. To confirm these findings, representative sections of native spleen, liver, aorta and heart were assessed and showed no gross morphological differences between groups (Supplementary Figure S5), suggesting no deleterious effects with the addition of PDE5i.

In the efficacy study, a significant reduction in tumour volume was observed in the CX, CX+Vardenafil and CX+Tadalafil (Two-way ANOVA, p-values of 0.003, <0.0001, and 0.007 respectively) with the biggest reduction in tumour volume in the CX+Vardenafil group. Comparison of the tumour volume in the 4 groups on the final day of the study revealed that only the CX+Vardenafil group was significantly reduced compared to vehicle control (Kruskal Wallis test, p=0.04), demonstrating the stronger effect of CX+Vardenafil compared to CX treatment alone or CX+Tadalafil.

## Discussion

Stromal remodeling can promote cancer progression. CAFs display an activated myofibroblast phenotype and expression of αSMA in many solid tumors is a marker of reduced disease free and overall survival^8, 35, 39-41^. The tumor-promoting biology of CAFs make them a target for novel cancer therapies. In this study, we have demonstrated that PDE5 is a potential new target for altering the fibroblast phenotype in EAC. Using a combination of conventional *in vitro* molecular biology techniques, state-of-the-art proteomic and single cell sequencing technologies we have documented the specificity of PDE5i for fibroblasts both to prevent transdifferentiation and revert the activated myofibroblast (CAF) phenotype. Finally, in a step towards a clinical trial we have confirmed the efficacy of PDE5i in combination with chemotherapy in close-to-patient *in vitro* and *in vivo* PDX-based model systems.

This study is not without shortcomings. Some of the *in vitro* work has been performed with cell lines that may not represent the true *in vivo* biology of these cell types. We supplemented these findings with those of more representative close-to-patient *in vitro* models which more accurately reflect the response of EAC cells to chemotherapy. Our proteomic analysis was conducted with a single, representative NOF/CAF pair, and further validation of these findings might be sensible. Finally, we have not tested PDE5i efficacy in a spontaneous EAC animal model. Unfortunately, no good model of EAC exists, meaning that the best testing-ground for PDE5i in EAC will be in humans. To mitigate this, we have used a validated near-patient PDX model to demonstrate efficacy.

Our findings are in keeping with many reports documenting the role of fibroblasts in cancer. Activated myofibroblasts are contractile and pro-invasive in EAC models^8^. There is evidence that CAFs can protect cancer cells from chemotherapy^42^, create an immunosuppressive environment, reduce the immune infiltrate and alter the immune composition allowing cancer cells to escape immune surveillance^43-45^. By reducing the trans-differentiation of fibroblasts in cancer and by modulating the phenotype of activated CAFs we may be able to improve overall survival by several different mechanisms: improving response to chemo/immuno-therapy, increasing tumor cell recognition by the immune system and reducing cancer cell invasion.

Several strategies have been proposed to target pro-tumorigenic CAF functions, mostly through modulating the effectors of the CAF phenotype rather than the cell state itself. These include targeting the ECM remodeling enzymes such as the lysyl oxidase family and MMPs, or targeting CAF-derived molecular signals (e.g. CXCR4, TGF-B, HGF) (reviewed in ^9^).

Initial attempts to specifically target CAFs have centered on the membrane bound glycoprotein Fibroblast Activation Protein alpha (FAP). Early promise with FAP-targeting monoclonal antibodies has not translated into clinical success (reviewed in^46^). Novel mechanisms to prevent myofibroblast differentiation and CAF accumulation are now required. PDE5i might offer a compelling way forward for this purpose. Importantly, PDE5i are a safe and well tolerated class of drug administered to millions of patients world-wide to treat erectile dysfunction, benign lower urinary tract symptoms and pulmonary arterial hypertension^21, 47^. High dose PDE5i show safety and efficacy for treating heart failure with reduced ejection fraction^48^. There is a significant body of evidence supporting PDE5i use in treating a range of cancers (reviewed in ^18^). In particular, animal studies suggest PDE5i have potent immunomodulatory activity that warrants clinical study with or without immune check-point inhibition^49^. PDE5i are currently being tested in combination with standard-of-care and other novel treatments in a range of cancer types including gliomas, head and neck squamous cell cancer, pancreatic cancer and malignant melanoma^18^. With this background, the sensible next steps for testing PDE5i in EAC are in the context of a phase I/II human clinical trial.

In summary, we provide *in vitro*, near-patient and *in vivo* evidence for the potential role of PDE5i in treating esophageal adenocarcinoma and suggest a rationale for future human trials.

---

**Supplementary Figure S1. A**. Western blot showing PDE5 siRNA knockdown in CAFs showing concomitant reduction in PDE5 and αSMA expression. **B**. Fibroblast contraction was assessed using CAFs ± PDE5 siRNA collagen-1 gel contraction assays. Reduction in PDE5 expression reduced collagen-1 gel contraction in CAFs and **(C)** the induction of FLO-1 cell invasion.

**Supplementary Figure S2. A**. Western blot and histogram assessing αSMA expression after 72 hours 50μM vardenafil treatment. A 50 % reduction in expression was achieved with one treatment in 72 hours, whereas treatment every 24 hours (3 treatments in a 72h period) significantly downregulated αSMA expression by 80%. **B**. Western blot assessing αSMA expression in different patient CAFs after the removal of vardenafil treatment. Both CAF populations underwent a significant reduction in αSMA, but comparison of the two CAF populations reveals differences in time to maximum inhibition and time to rebound levels of αSMA expression.

**Supplementary Material S3. Further details on chemosensitivity findings in 3D-TGA cultures**.

**Supplementary Figure S4. PDE5i dose escalation study in combination with ECX at 50% dosage in non-tumor-bearing mice**. Immune-incompetent mice (n=3) tolerate ECX treatment at half the equivalent human dosage. 3 cycles of ECX were administered over 8 weeks. Cycle 1 was 25% equivalent dose; Cycle 2 was 50% equivalent dose; In cycle 3, two mice received 75% equivalent dose and one mouse received a repeat of cycle 2 due to rapid weight loss (50%).

**Supplementary Figure S5. Histological evaluation of PDX tumors and murine organs previously reported as being affected by ECX treatment**. Hematoxylin & eosin stained sections of murine tissues from PDX-bearing mice in this study, treated with ECX alone or ECX with vardenafil or tadalafil. No gross histological changes were observed in mice treated with vardenafil or tadalafil compared to mice receiving ECX treatment alone.

## Notes

**Conflicts of interest:** The authors declare no conflict of interest.

**Funding:** Medical Research Council UK Clinician Scientist Fellowship to T.J.U. “Exploring stromal-epithelial interactions in oesophageal cancer” G1002565 IDN0. 99762; Cancer Research UK & Royal College of Surgeons of England Advanced Clinician Scientist Fellowship to T.J.U. “Cellular interplay in oesophageal cancer.” A23924.

### Competing Interest Statement

The authors have declared no competing interest.

