## Supplementary Methods for "Modulation of the tumour promoting functions of cancer associated fibroblasts by phosphodiesterase type 5 inhibition increases the efficacy of chemotherapy in human preclinical models of esophageal adenocarcinoma"

**Cell lines**

FLO-1 (EACC), OE33 (EACC), MFD-1 and primary cells were maintained in Dulbecco’s modified Eagles medium (DMEM, Invitrogen) or Roswell Park Memorial Institute medium (RPMI; Invitrogen) supplemented with 10% (v/v) fetal calf serum (FCS, Autogen Bioclear), 2 mM L-glutamine and 100 μg/mL penicillin/streptomycin (Invitrogen). The MFD-1 cell line is a well characterized primary esophageal cancer cell line and has been described previously^1^.

**SiRNA transfections**

Commercially available PDE5 sequences 1, 5’-CCAGUGCUCAAGACUCUUGtt-3’, 3’-ctGGUCACGAGUUCUGAGAAC -5’ (137131, Ambion) and Sequence 2, 5’-GCAAGCUAUUUUAGCUACAtt-3’, 3’-ttCGUUCGAUAAAAUCGAUGU-5’ (137133, Ambion) were transfected into primary fibroblasts at 50% confluence (cells were seeded 24 hours before transfection, 5x10^5^/well of a 6 well plate) using INTERFERin (Polyplus), according to the manufacturer’s instructions. Cells were cultured for 72 hours to achieve optimal knockdown, negative control 1 siRNA (Ambion) was used for experimental controls.

**Western blotting**

Antibodies used: mouse monoclonal anti-α-SMA (M085129-2, Dako), mouse monoclonal anti-HSC-70 (sc-7298, Santa Cruz) and rabbit polyclonal anti-PDE5 (SC-32884, Santa Cruz). Normal Oesophageal fibroblasts were pre-treated with 50 μM vardenafil (Sigma) for 1 hour before treatment with recombinant TGF-β1 (Calbiochem) for 72 hours. Cancer associated fibroblasts were treated with 50 μM vardenafil for 72 hours or 3x 50 μM vardenafil over 72 hours. Cells were harvested by trypsin digestion after an initial PBS wash before pelleting by centrifugation. Cell lysis was carried out for 15 minutes at 4°C in 50μl RIPA buffer (0.75M NaCl, 5% NP40, 2.5% deoxycholic acid, 0.5% SDS, 0.25M Tris pH8.0). Lysates were clarified by centrifugation at 8000xg for 5 minutes. Fibroblasts or FLO-1 cells were cultured in serum free DMEM for 24 hours before the conditioned medium was harvested at 4°C, clarified by centrifugation and the supernatant concentrated using Amicon ultra-4-centrifugal 10kDa filters (Millipore). Protein samples were quantified using the Bradford protein assay reagent. 20 μg of protein or 20μl of concentrated cell culture medium (adjusted in SDS loading dye for cell number when conditioned medium was removed) were resolved using SDS-polyacrylamide electrophoresis and transferred to Hybond-ECL membranes (GE healthcare). Blocking and antibody incubations were done in 3% low-fat milk in PBS–0.025% Tween 20, and washes were in PBS–0.1% Tween 20. Detection of horseradish peroxidase-labelled secondary antibody was done with Supersignal (Pierce), and images were collected using a CCD camera (ChemiDoc-it® imaging system, UVP).

**Proteomics**

*Database searching*

Unprocessed raw files were submitted to Proteome Discoverer 1.4 for target decoy search using the Sequest algorithm as reported previously^2, 3^. The UniProtKB homo sapiens database which comprised 20,159 entries (release date January 2015) was utilized. FDR corrected *p-value* at the peptide level was set at <0.05. Percent co-isolation excluding peptides from quantitation was set at 50. Reporter ion ratios from unique peptides only were taken into consideration for the quantitation of the respective protein.

The iTRAQ ratios of proteins were median-normalized and log_2_ transformed. Principal component analysis of all quantified proteins was performed using ClustVis (<https://biit.cs.ut.ee/clustvis/>). A one-sample Student’s T-Test was performed to identify differentially expressed proteins (DEPs) in treated cells vs. their respective controls as one group. Proteins identified with at least two unique peptides and a one-sample Student’s T-Test (*p-value*<0.05) were considered differentially expressed. Gene ontology analysis was performed using DAVID (<https://david.ncifcrf.gov/summary.jsp>). A Fisher-exact p-value < 0.05 was considered significant.

**Droplet barcoded single cell RNA sequencing**

Captured mRNA was reverse transcribed and the resulting cDNA libraries were amplified, purified and prepared for sequencing using a modified Nextera XT protocol and sequenced using Illumina NextSeq500. Sequenced reads were aligned to Genome Reference Consortium Human Build 37 (HG19) and processed using DropSeq tools 1.12 to produce a digital expression matrix where columns are cells and rows are genes, gene counts were created by counting UMIs. Clustering and differential expression analysis was performed using R version 3.5.1 (Feather Spray) and the package Seurat version 2.3.4.

Raw data consisted of 1800 cells expressing 24,434 genes. After removing genes expressed in less than 5 cells and cells with less than 1500 genes we created a Seurat object with 1122 cells expressing 18,436 genes. Gene counts were log normalised and scaled to read depth and mitotic phase to remove unwanted sources of variation using Seurat’s CellCycleScoring and ScaleData functions.

The 3,554 most highly variable genes (highest log variance to mean ratio and highest mean expression) were used to perform principal component(PC) analysis. The first principal component consisted of genes known to be markers of fibroblasts and fibroblast activity, such as Decorin, vimentin and CD90 (THY1) and genes known to be markers of adenocarcinoma such as EPCAM and the cytokeratins.

The first 10 principal components were used as input to Seurat’s FindClusters function and dimensionality reduction was performed using RunTSNE. This first pass clustering produced some populations from the co-cultured cells that were expressing known markers of fibroblasts and cancer cells suggesting that they were doublets (two cells exposed to one nanobead). We identified these cells by highlighting all cells that had a scaled expression of greater than one for any of the top 30 fibroblast markers from PC 1 and any of the top 30 markers for cancer cells from the same PC. We removed 109 cells from our dataset 104 from the co-culture experiments. We then repeated the clustering steps above without the “doublet” population.

**3D-tumor growth assay**

The full method for the establishment of close-to-patient OAC cells using a feeder layer method and subsequent growth with a stromal component in the 3D-tumour growth assay to form cancer cell clusters, which can then undergo clinically relevant *ex vivo* pharmacological assessment has been published by our group^1^.

In this *ex vivo* study we undertook evaluation of novel adjunct PDE5i administration in additional to the regimen of Epirubicin, Cisplatin and 5-Fluorouracil / Capecitabine. This is the standard of care pre and post-operative chemotherapy regimen used for the treatment of OAC in the UK, and was administered to patients clinically in this study. This regimen was replicated in the 3D-TGA, with and without the stromal component of the assay, and with and without the adjunctive PDE5i Vardenafil. The chemotherapeutic effect of the drugs evaluated in the 3D-TGA is calculated as a percentage of the matched untreated control. Using the Chou Talalay method^5^ IC50 curves are generated for the drugs both individually and in combination as previously described by our group^4^. The IC50 values were compared with the mean peak serum concentrations seen in patients for each of the drugs (see table below). The chemotherapeutic response was thus defined as sensitive (IC50 below the mean peak serum), borderline (+/- 10% of the mean peak serum), or resistant (IC50 above the mean peak serum).

Statistical analysis was undertaken to assess the relative efficacy of the drug combinations using a two-way ANOVA to compare the different parameters among the different groups. Difference between groups was only considered to be significant if there was no overlap between the 95% confidence interval about the median. The paired and un-paired t-test was used to calculate the significance of difference between parametrically distributed groups, and Mann-Whitney U test between independent groups, with a significance level of p < 0.05. Statistics were computed with GraphPad Prism Software (San Diego, CA, USA) and plotted with mean values, and error bars for standard deviation.

Peak serum concentration of chemotherapy agents in humans

| **Drug** | **Mean peak serum** | **Reported publications** | **Dose in reference** |
| --- | --- | --- | --- |
| **Epirubicin** | 4.5µM | ^5-9^ | 50mg/m^2^ |
| **Cisplatin** | 4.3µM | ^10-12^ | 60 mg/m^2^ |
| **Fluorouracil / Capecitabine** | 4.6µM | ^13-15^ | 200 mg/m^2^  & 625 mg/m^2^ |
| **Vardenafil** | 0.0267μM | ^18^ | 20mg for ED |

**Patient derived xenograft models**

The *in vivo* experiments were conducted under the UK Home Office Licence number PPL P435A9CF8. LASA good practice guidelines, FELASA working group on pain and distress guidelines and ARRIVE reporting guidelines were also followed.

All mice were purchased from Charles River UK. Mice were maintained in individually Ventilated Cages (Tecniplast UK) within a barriered unit, illuminated by fluorescent lights set to give a 12 hour light-dark cycle (on 07.00, off 19.00), as recommended in the guidelines to the Home Office Animals (Scientific Procedures) Act 1986 (UK). The room was air-conditioned by a system designed to maintain an air temperature range of 21 ± 2ºC and a humidity of 55% ± 10%. Mice were housed in social groups, 3 per cage, during the study, with irradiated bedding and autoclaved nesting materials and environmental enrichment (Datesand UK). Sterile irradiated 5V5R rodent diet (IPS Ltd, UK) and irradiated water (Baxter, UK) was offered *ad libitum.* The condition of the animals was monitored throughout the study by an experienced animal technician. After a week’s acclimatisation, the mice were initiated with tumours as described below.

**Dose escalation study**

The dose escalation was performed as follows:

10 male 8-9 week old CD-1 NuNu mice were implanted with 1x10^6^ OES127 cells re-suspended in 100μl of Matrigel (Corning), which were developed from an esophageal adenocarcinoma resection specimen + eGFP labelled mesenchymal stem cells (MSCs) in a ratio of 2:1. The cells were generated by disaggregating from donor OES127 PDXs by a collagenase/dispase disaggregation fluid, and rotating at 37C for 1 hour, counted and viability measured by trypan blue, before resuspending both the MSCs and PDX simultaneously in matrigel. These were injected subcutaneously into the left flank of the mice and the resulting tumours were measured twice weekly using Vernier calipers and the volumes calculated using the formula V=ab2/6, where a is the length and b is the width. A secondary dose of MSCs, this time lentivirally transduced with pLVX-fLuc, were added as a ‘boost’, 14 days after initiation, directly injected into the tumour in Phosphate Buffered Saline (Sigma, UK). Dosing commenced on day 18 post-initiation and followed the regime below, with a 14 day rest period between each cycle for observation of side effects, during which time animals were weighed daily.

Dosing regime

- Group 1 Control-no treatment
- Group 2 Epirubicin 15mg/kg, IV, day 1 + Cisplatin 3mg/kg, IP, days 1, 3, 5 + Capecitabine 100mg/kg, PO, days 1, 2, 3, 4, 5 (twice daily) [ECX]
- Group 3 ECX (as above) plus Vardenafil, twice daily at:
  - 2mg/kg (i.e. 4mg/kg/day), PO, days 1, 2, 3, 4, 5
  - 6mg/kg (i.e. 12mg/kg/day), PO, days 15, 16, 17, 18, 19, 20
  - 8mg/kg (i.e. 16mg/kg/day), PO; days 29, 30, 31, 32, 33
- Group 4 ECX (as above) plus Tadalafil, twice daily at:
  - 2mg/kg, (i.e. 4mg/kg/day), PO, days 1, 2, 3, 4, 5
  - 4mg/kg, (i.e. 8mg/kg/day), PO, days 15, 16, 17, 18, 19, 20
  - 10mg/kg, (i.e. 20mg/kg/day), PO, days 29, 30, 31, 32, 33

*IV=intravenous, IP=intraperitoneal, PO=per os, SFI=saline for injection, WFI= water for injection

As this was a dose escalation tolerability study, no power calculation was required, groups 1 and 2 having 2 mice each and groups 3 and 4 having 3 mice each. The mice were terminated between days 42-56 due to tumours approaching maximum allowable size. They were culled by cervical dislocation, tumours were dissected out and weighed, aortas, hearts, livers and spleens were also dissected out, and all were fixed in Neutral Buffered Formalin.

Due to some slight weight loss in one group (ECX) during this dose escalation study which was thought to be associated with the use of epirubicin, an additional tolerability study was carried out using cisplatin and capecitabine only, at 75% of those used in the Dose Escalation and using the PDE5is  daily (instead of twice daily at lower doses). This dosing regime, which was well-tolerated was adopted for the Efficacy study (see details below).

**Efficacy study**

The efficacy study was performed as follows:

60 female x 7-8 week old CD-1 NuNu mice were implanted with OES127 cells as described above. Tumours were measured as previously detailed, and mice were weighed weekly. Tumours were also imaged weekly in the IVIS® Spectrum imaging system (PerkinElmer, MA, USA) by 2D optical imaging, with tumour measurements made using Living Image (4.3.1) software and standard open filters to assess the retention of MSCs.

Prior to imaging, the mice were anaesthetised with an injectable anaesthetic combination (Anaestemine [ketamine]/Sedastart [medetomadine], Animalcare Ltd. UK) before being placed in the IVIS system and imaged on days 0, 1, 8, 15, 16, 21, 23 and 28, mice being allowed to recover from the anaesthetic with appropriate post procedural monitoring and therapy, including placing mice on a heat pad and providing fluid replacement via wet mash once awake.

On Day 14 after tumour initiation, the mice were randomised into group of 15 mice per group by tumour size and fluorescence. Power calculations to determine group sizes were based on One way ANOVA with 4 groups, to allow detection of a 40% effect of treatment at a power of 80%. This gave a minimum required sample size of n=10 per group but based on a potential 70% take rate (due to the tissue being PDX in origin and the implant location) sample size was increased to n=15 per group. Dosing followed the weekly cycle below for 3 weeks, with 2 days dosing in week 4 prior to termination on week after final dosing.

Dosing regime

- Grp 1 Cisplatin 2.25mg/kg, IP, days 1, 3, 5 + Capecitabine 75mg/kg, PO, days 1, 2, 3, 4, 5 [CX]
- Grp 2 CX (as above) plus Vardenafil (16mg/kg, PO) days 1, 2, 3, 4, 5
- Grp 3 CX (as above) plus Tadalafil (20mg/kg, PO) days 1, 2, 3, 4, 5
- Grp 4 Placebo dosed control, SFI vehicle IP days 1, 3, 5 and WFI vehicle PO days 1, 2, 3, 4, 5

The mice were culled by cervical dislocation, tumours were dissected out and weighed, before half was snap frozen in liquid nitrogen, half was fixed in Neutral Buffered Formalin. Data from one of the mice in Group 2 was excluded from the final analysis because it reached the maximum allowed size a week before any of the other mice in the study needed to be terminated, and thus was considered an outlier. Growth of tumours was assessed based on calliper measurements and expressed as a percentage of the pre-treatment volume for individual mice. Mean and standard error was calculated for each group and analysed by 2-way ANOVA to compare each group to the untreated group and by Kruskal-Wallis test to compare relative tumour volume between the groups at the final timepoint.

18.	Bayer. Levitra (Vardenafil hydrochloride) Product scientific and clinical information. 2014.
