## Supplementary figures and images for "Modulation of the tumour promoting functions of cancer associated fibroblasts by phosphodiesterase type 5 inhibition increases the efficacy of chemotherapy in human preclinical models of esophageal adenocarcinoma"

### Supplementary Figure S1

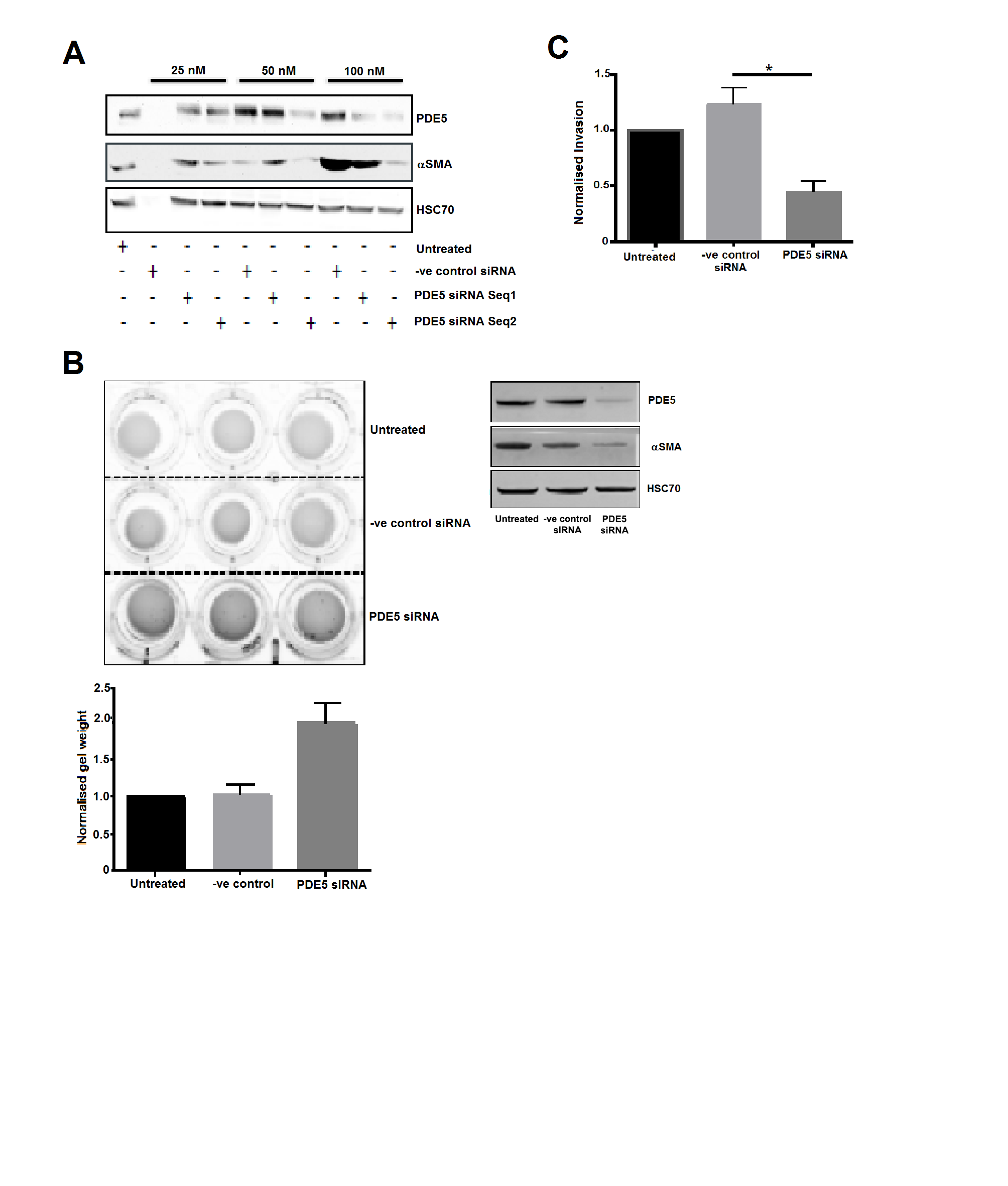

### Supplementary Figure S2

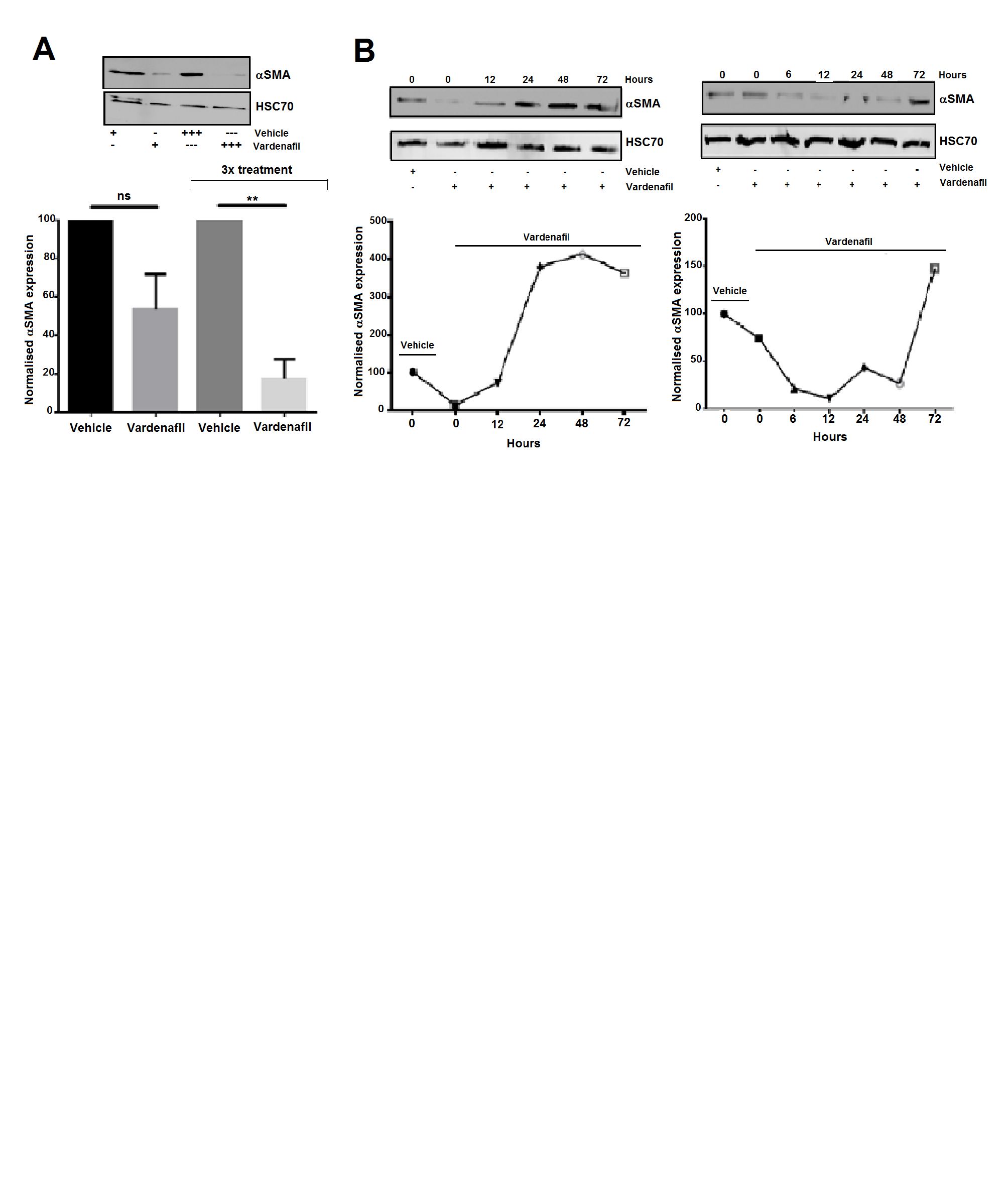

### Supplementary Figure S4

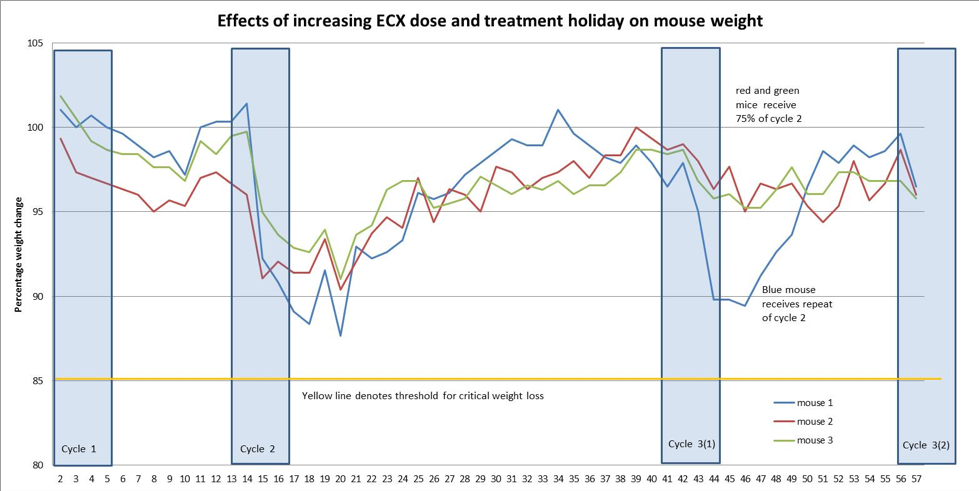
