## Supplementary Material S3 for "Modulation of the tumour promoting functions of cancer associated fibroblasts by phosphodiesterase type 5 inhibition increases the efficacy of chemotherapy in human preclinical models of esophageal adenocarcinoma"

**Supplementary Material – 4**

Figure 1

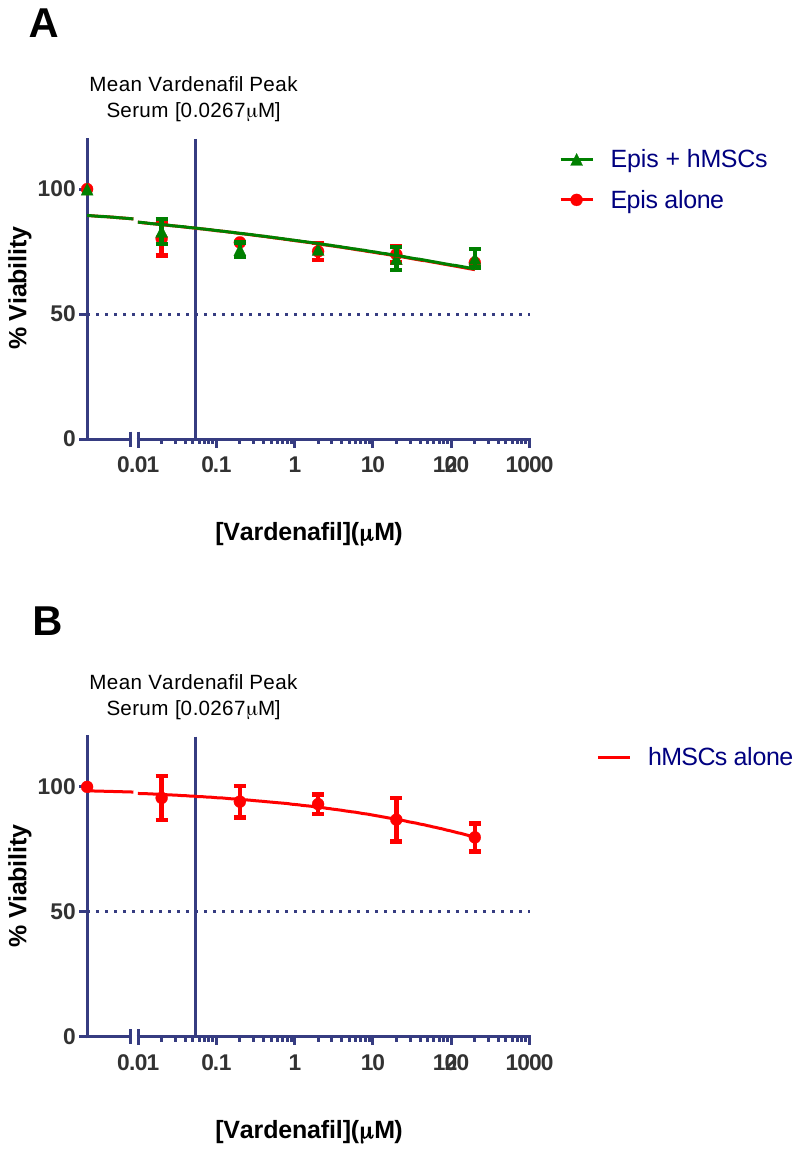

Figure 1. Cells grown in the 3D-TGA do not demonstrate chemo-toxicity upon exposure to the PDE5i Vardenafil at human relevant concentrations

The chemo-sensitivity of oesophageal adenocarcinoma (OAC) cancer cell clusters to Vardenafil was determined in 3D-TGA, in 6 replicate wells, after 4 day exposure to the drug at a range of concentrations, using the alamarBlue assay to measure viability. Viability curves were generated and IC_50_ values calculated using GraphPad Prism. Error bars represent one standard deviation. **(A)** OAC epithelial cells are grown in the 3D-TGA with and without hMSC co-culture before exposure to increasing concentrations of Vardenafil. **(B)** hMSCs are grown without epithelial cancer cells in the 3D-TGA before exposure to increasing concentrations of Vardenafil.

Figure 2

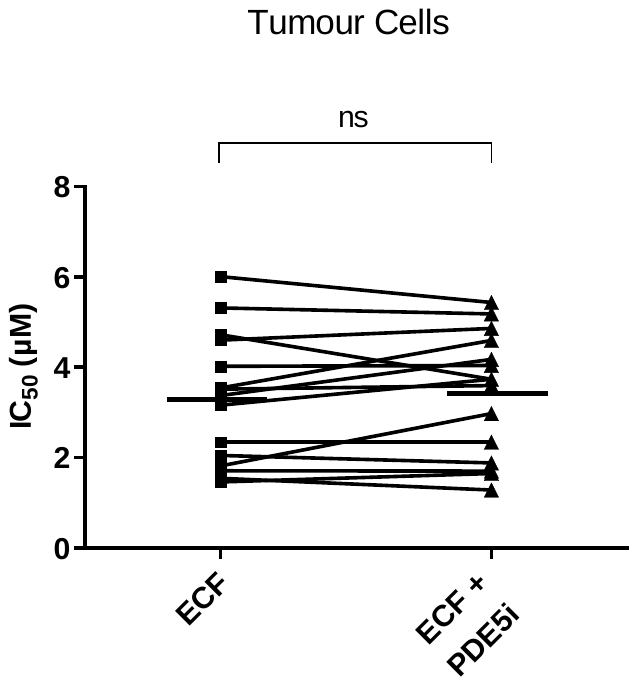

**Figure 2. In the absence of hMSC co-culture, combination therapy of ECF chemotherapy with the PDE5i vardenafil does not provide enhanced chemo-sensitivity**

Sensitivity of close-to-patient cells was determined in 3D-TGA, without mesenchymal cell co-culture, after 4 day exposure to ECF and Vardenafil (PDE5i) drug combinations. Viability curves were generated and IC_50_ values determined using GraphPad Prism. Horizontal lines represent mean IC_50_s.

Overall sensitivity of all of the patient cell lines cultured as Tumour cells alone (**A**) was determined for assays with and without the addition of PDE5i to ECF chemotherapy. Statistical significance calculated using the paired t-test shows no difference between the two groups.

Figure 3 and Figure 4

The effect of PDE5i addition is different on individual patient OAC cells which demonstrate hMSC co-culture induced chemo-resistance to ECX chemotherapy.

The OAC sample Oes5B(ii) is used as an example to demonstrate the effect for an individual patient of addition of the PDE5i drug Vardenafil to the standard ECF chemotherapy.

This patient’s *ex vivo* 3D model demonstrates a significant mesenchymal co-culture-induced ECF chemotherapy resistance (Figure 3A).

The PDE5i addition to ECF chemotherapy exhibits an enhanced chemo-toxic effect (Figure 3B), which does not occur in the absence of stromal co-culture (Figure 3C) suggesting that the PDE5i drug is acting upon the stromal component of the 3D-TGA co-culture model.

This patient’s cancer cells are resistant to treatment with conventional ECF chemotherapy clinically (with a TRG of 4) and also chemo-resistant in the 3D model with stromal co-culture, with an *ex vivo* an IC50 of 4.977 (Figure 3D). However, when the patient’s co-cultured cancer is treated with the PDE5i drug Vardenafil in addition to the standard ECF chemotherapy, the cancer cells were found to be sensitive to the chemotherapy, at a clinically-relevant dose with an IC50 of 2.105 (Figure 3D).

In contrast, although the individual patient OAC Oes6B(i) also demonstrates a significant mesenchymal co-culture induced ECF chemotherapy resistance (Figure 4A), the addition of PDE5i does not exhibit an additional chemo-toxic effect and the 3D model continues to exhibit stromal induced chemo-resistance (Figure 4B).

Therefore in this patient’s co-culture model, addition of PDE5i to the ECF chemotherapy does not have any effect on chemo-sensitivity (Figure 4D), and the IC50 remains greater than the mean peak serum clinically administered to patients and resistant to ECF chemotherapy, as it is clinically with a TRG of 4.

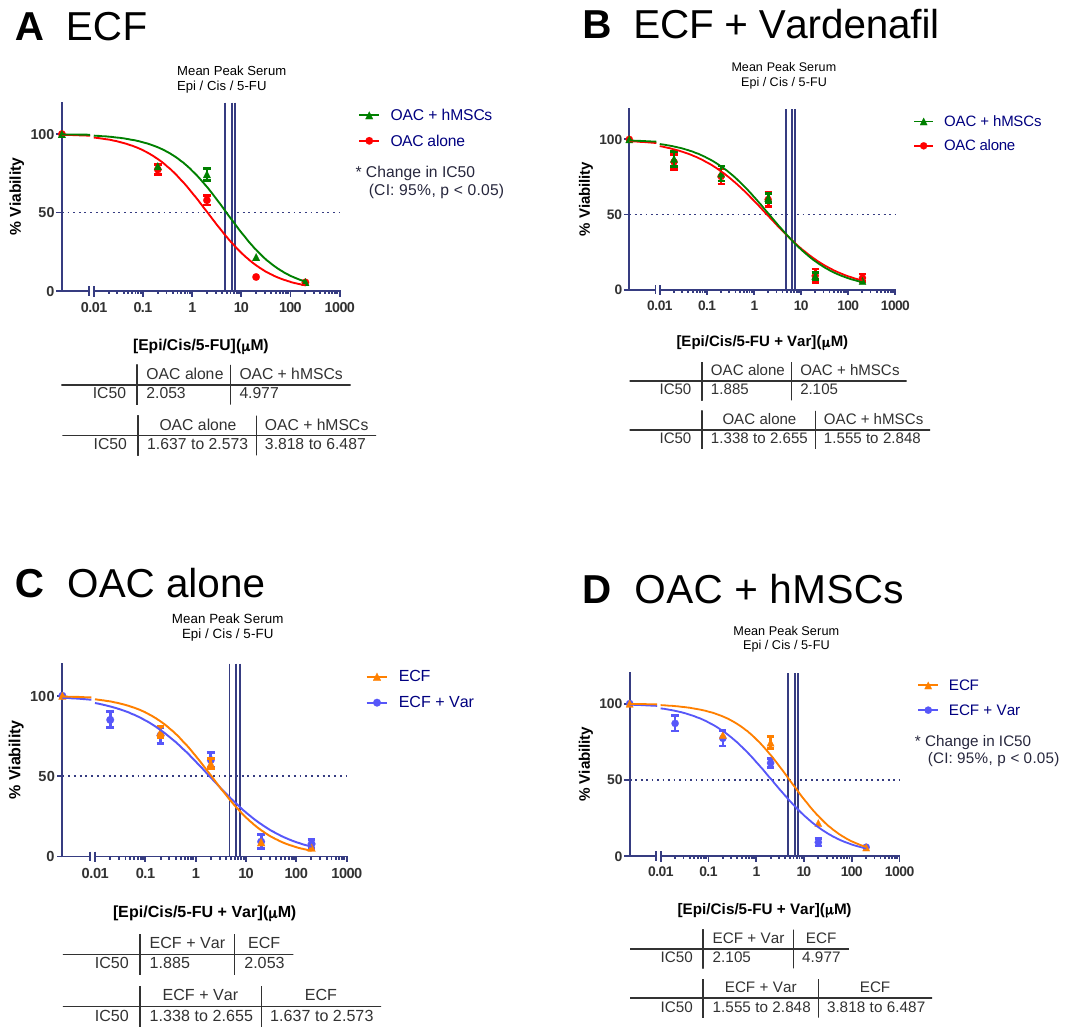

Figure 3. The alternate effects of PDE5i on close-to-patient OAC cells demonstrating hMSC co-culture induced chemo-resistance to ECX chemotherapy: *Addition of the PDE5i agent Vardenafil can increase chemo-sensitivity.*

Close-to-patient OAC cells are cultured in the 3D-TGA with and without hMSC co-culture, before exposure to the standard of care chemotherapy agents Epirubicin, Cisplatin and Fluorouracil with and without the presence of the PDE5i drug Vardenafil.

(**A**) Resistance to ECX chemotherapy is significantly increased with hMSC co-culture. (**B**) Increased chemo-resistance with hMSC co-culture is no longer observed by the addition of PDE5i to ECX chemotherapy. (**C**) When epithelial cells are grown alone without hMSC co-culture, no change in chemotherapy sensitivity is seen by the addition of PDE5i to ECX chemotherapy. (**D**) When epithelial cells are co-cultured with hMSCs, addition of PDE5i to ECX chemotherapy results in a significant increase in chemo-sensitivity.

Statistical significance is determined by the arithmetic variance between the calculated IC_50_s and their distinct 95% confidence intervals which have no overlap.

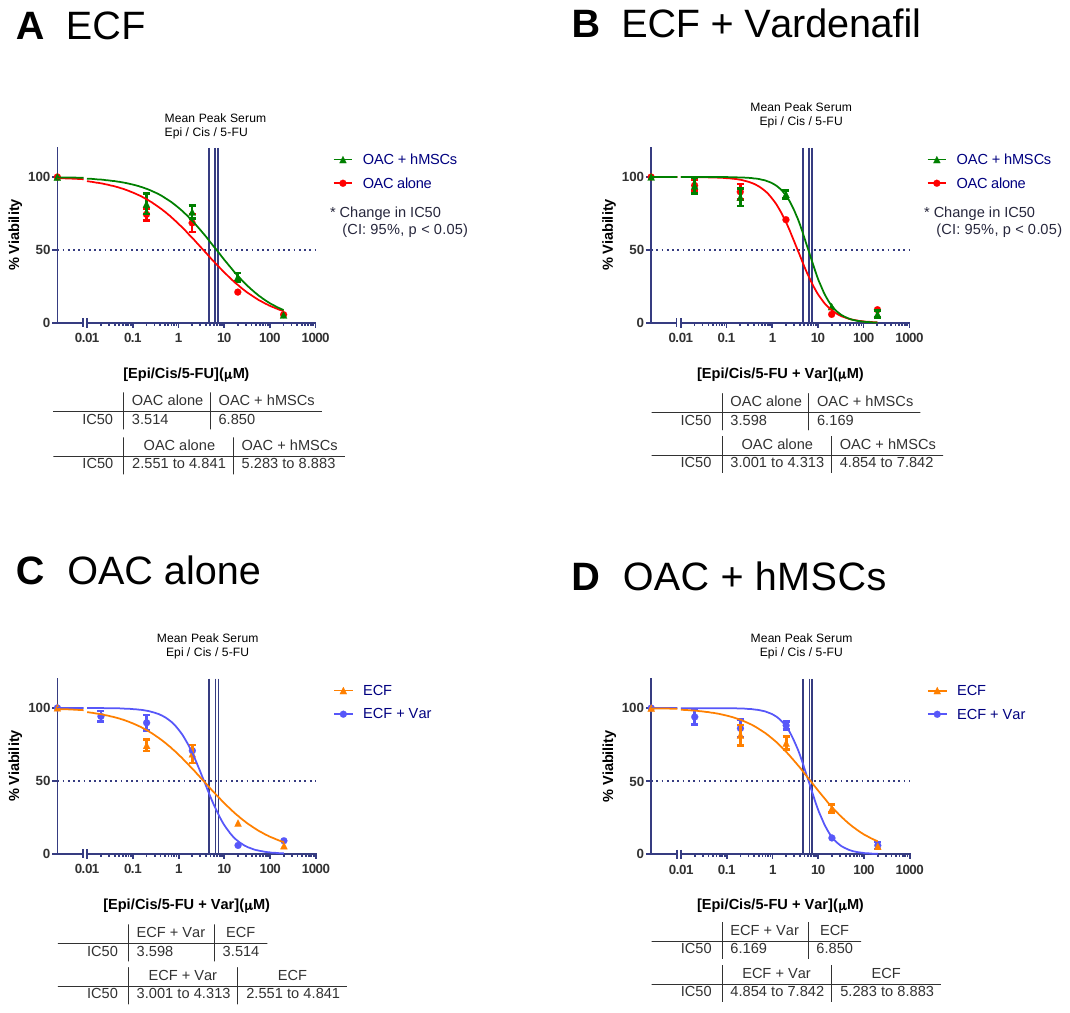

Figure 4. The alternate effects of PDE5i on close-to-patient cells demonstrating hMSC co-culture induced chemo-resistance to ECX chemotherapy: *Addition of the PDE5i agent Vardenafil can have no effect upon chemo-sensitivity.*

Close-to-patient OAC cells are cultured in the 3D-TGA with and without hMSC co-culture, before exposure to the standard of care chemotherapy agents Epirubicin, Cisplatin and Fluorouracil with and without the presence of the PDE5i drug Vardenafil.

(**A**) Resistance to ECX chemotherapy is significantly increased with hMSC co-culture. (**B**) Increased chemo-resistance with hMSC co-culture remains significant and is not affected by the addition of PDE5i to ECX chemotherapy. (**C**) When epithelial cells are grown alone without hMSC co-culture, no change in chemotherapy sensitivity is seen by the addition of PDE5i to ECX chemotherapy. (**D**) When epithelial cells are co-cultured with hMSCs, addition of PDE5i to ECX chemotherapy does not have any effect on chemo-sensitivity.

Statistical significance is determined by the arithmetic variance between the calculated IC_50_s and their distinct 95% confidence intervals which have no overlap.
