## Supplementary Figure S5 for "Modulation of the tumour promoting functions of cancer associated fibroblasts by phosphodiesterase type 5 inhibition increases the efficacy of chemotherapy in human preclinical models of esophageal adenocarcinoma"

| Table of Histology: CSU 1651c  Comparison of murine organs previously reported as histologically affected by administration of high dose PDE5i, demonstrates no histological change between the baseline Epirubicin/Cisplatin/Capecitabine (ECX) chemotherapy regimen and addition of PDE5i drugs | | | |
| --- | --- | --- | --- |
| Tissue | ECX alone | ECX + Vardenafil | ECX + Tadalafil |
| Snap-shot of Tumour | 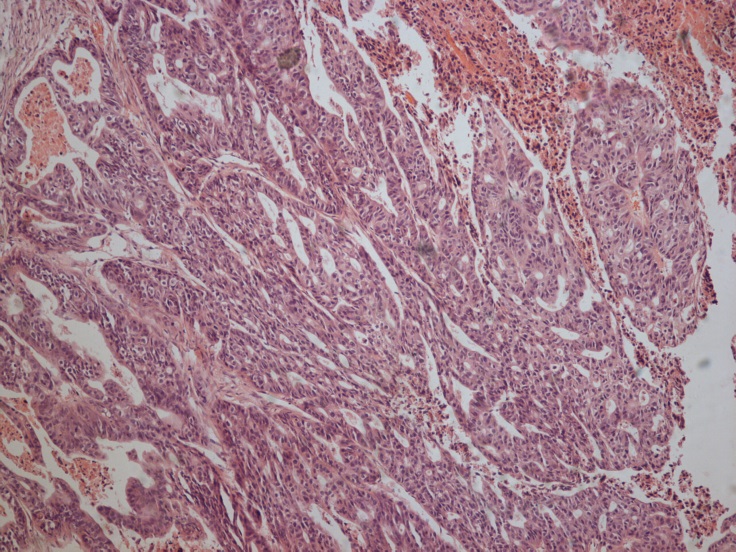 | 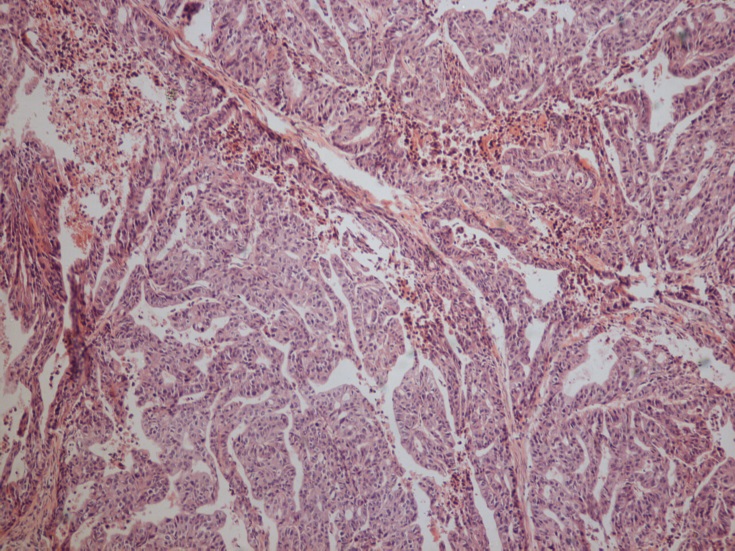 | 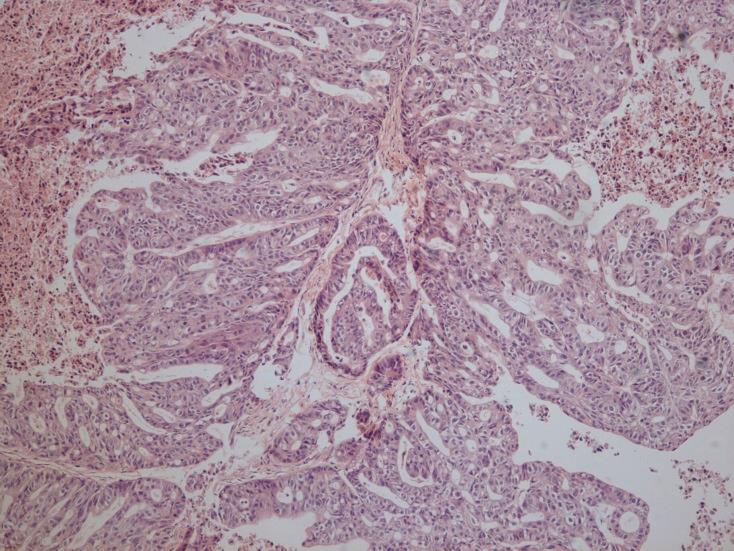 |
| Spleen | 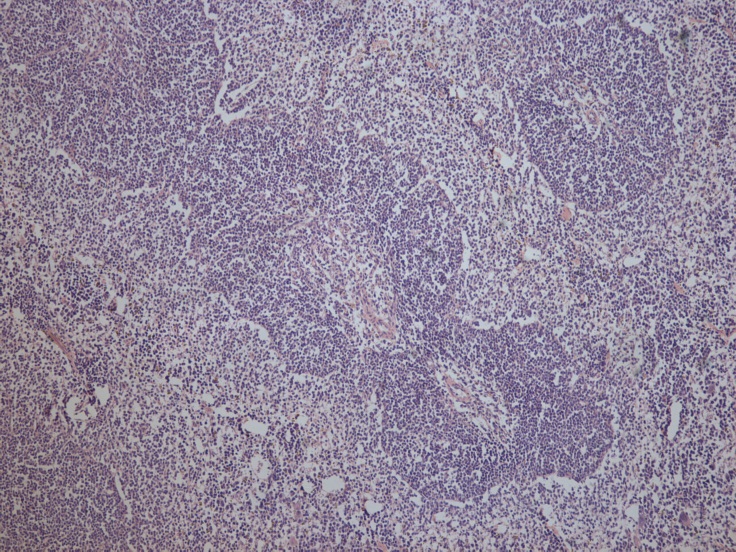 | 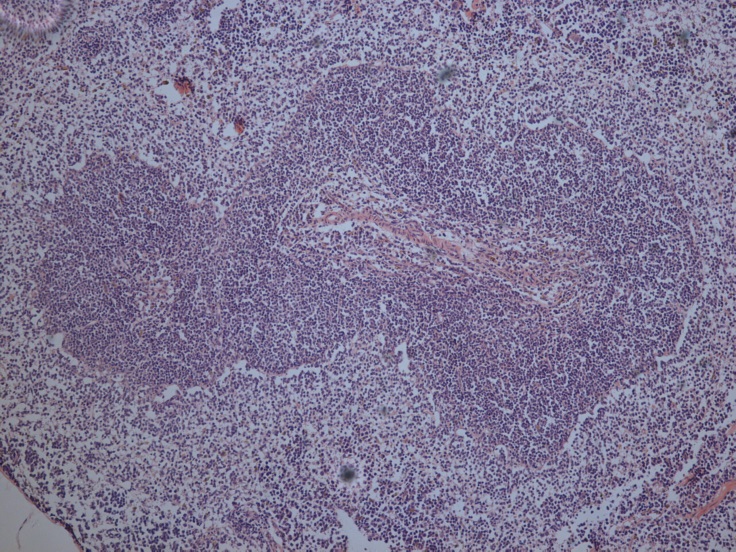 | 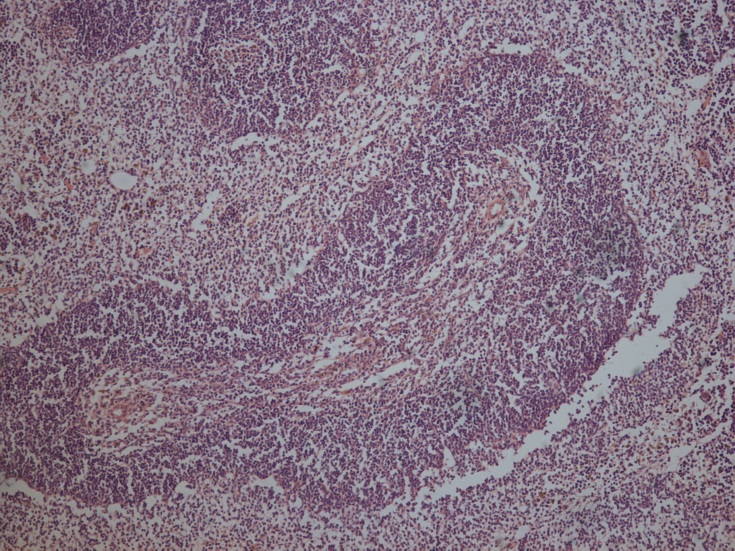 |

| Liver | 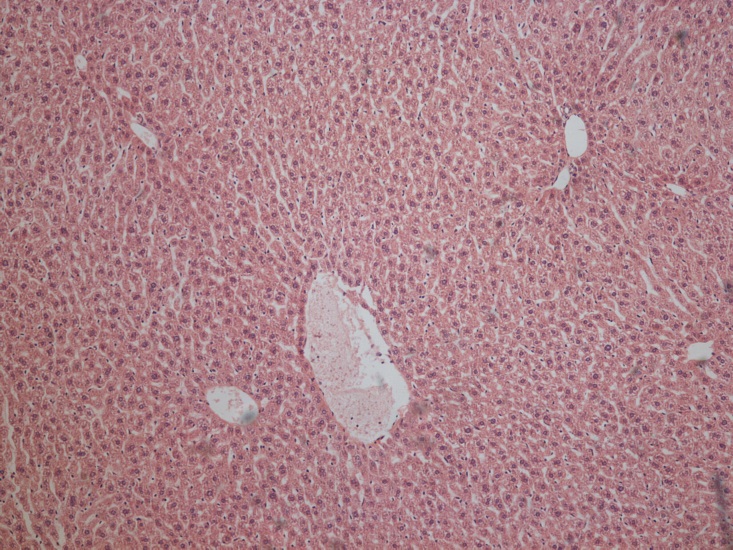 | 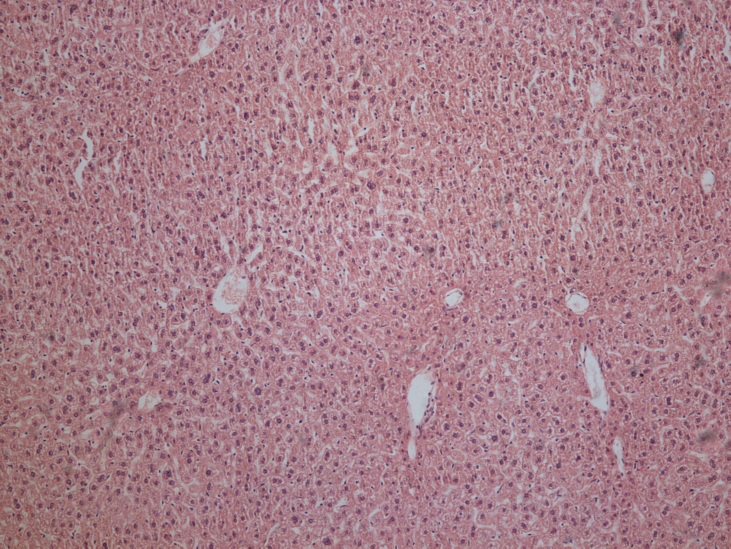 | 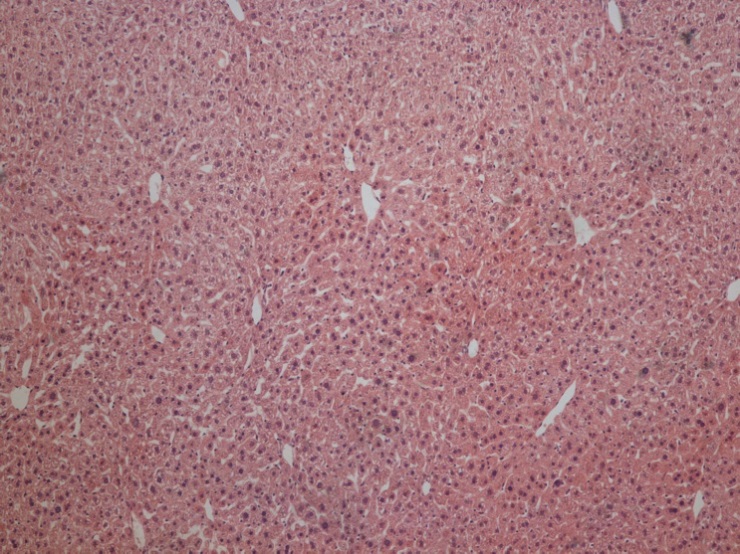 |
| --- | --- | --- | --- |
| Heart | 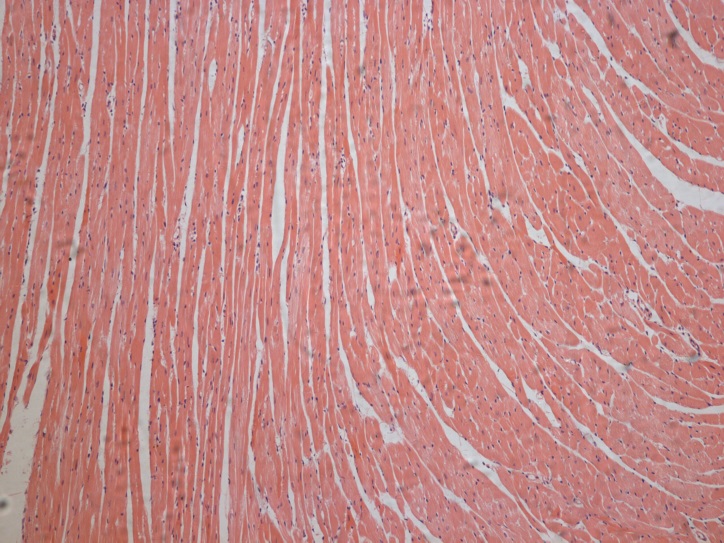 | 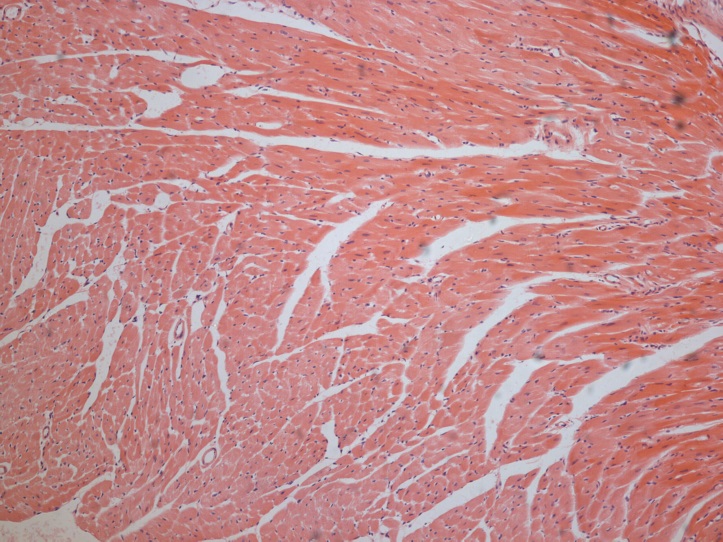 | 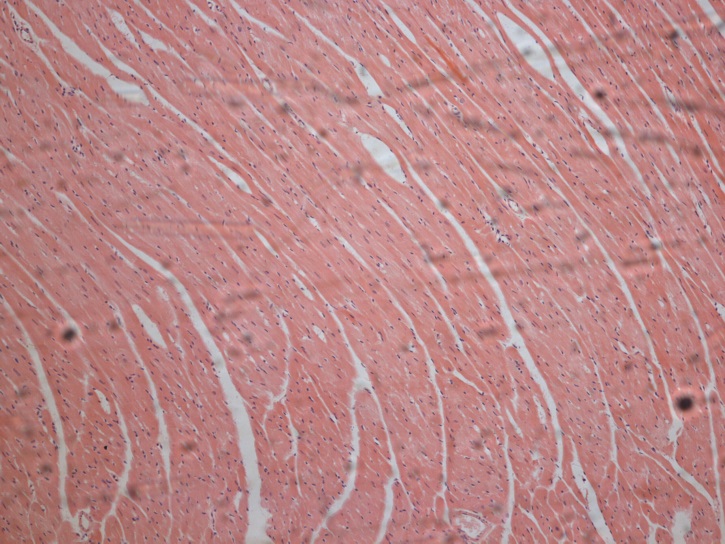 |

| Aorta (x10) | 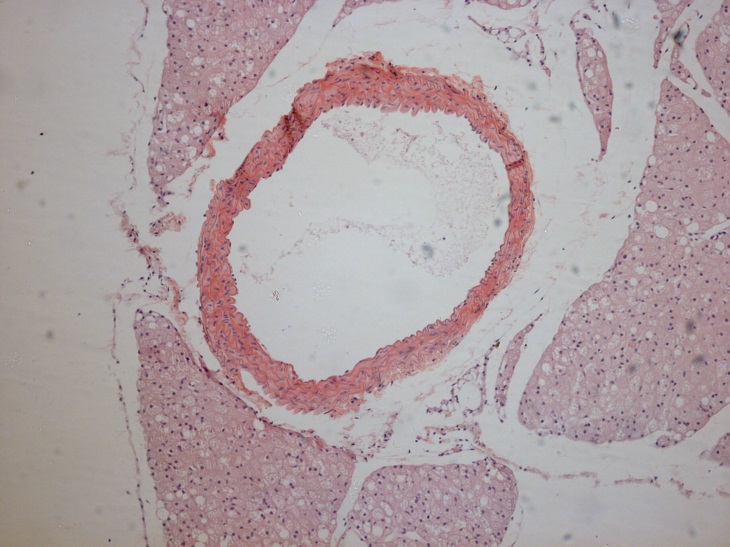 | 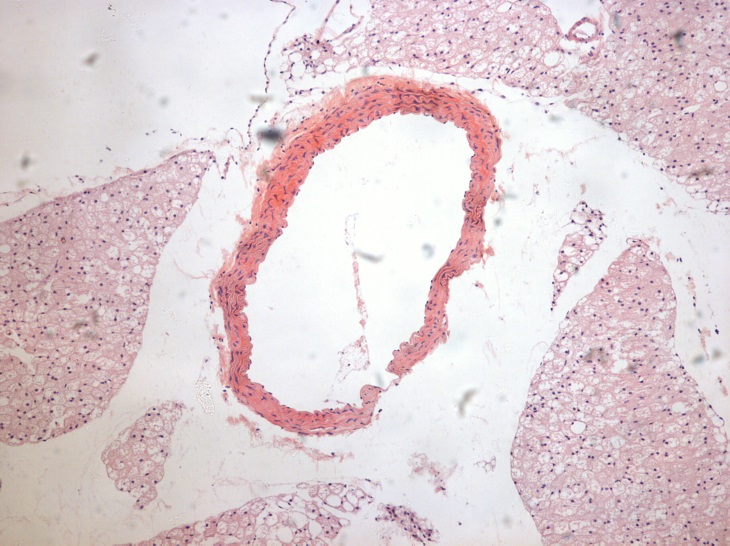 | 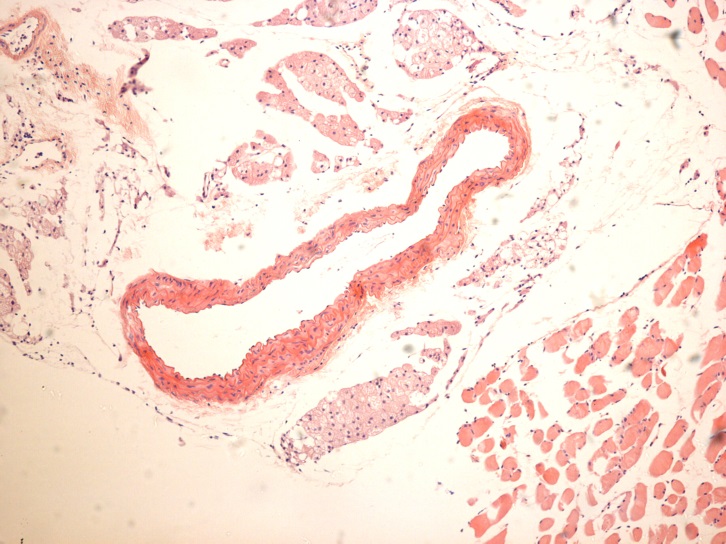 |
| --- | --- | --- | --- |
| Aorta (x40) | 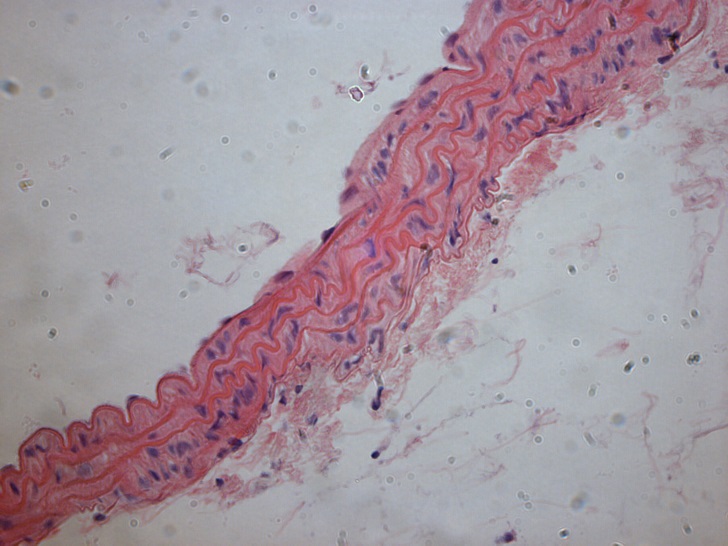 | 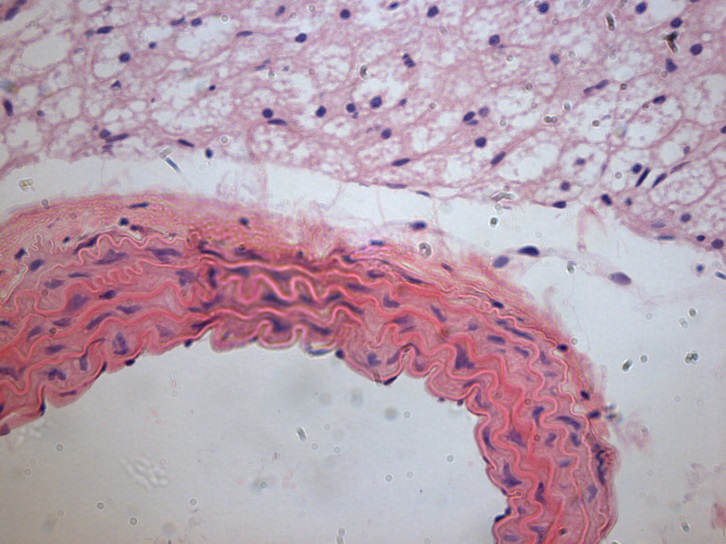 | 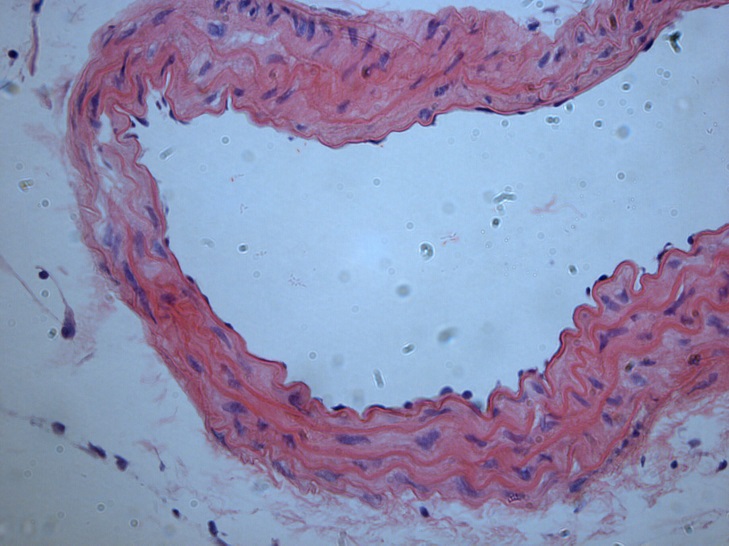 |
